# Iterative and modular expression of *Botryococcus braunii* genes enhances isoprenoid production in the diatom *Phaeodactylum tricornutum*

**DOI:** 10.1101/2025.11.27.690994

**Authors:** Luca Morelli, Cecilie Jensen, Eva C. Arnspang, Michele Fabris

**Affiliations:** SDU Biotechnology, Department of Green Technology, Faculty of Engineering, University of Southern Denmark, Campusvej 55, Odense M, DK-5230, Denmark; SDU Climate Cluster, Faculty of Science, University of Southern Denmark, Campusvej 55, Odense M, DK-5230, Denmark

**Keywords:** *Botryococcus braunii*, *Phaeodactylum tricornutum*, carotenoids, triterpenoids, squalene, synthetic biology, DBTL

## Abstract

Isoprenoids are essential natural compounds with high structural and functional diversity. Among them, carotenoids and triterpenoids such as squalene have high biotechnological value but remain challenging to produce sustainably. The green microalga *Botryococcus braunii* synthesizes large amounts of triterpenoids through specialized methylerythritol phosphate (MEP) pathway configurations involving distinct 1-deoxy-D-xylulose-5-phosphate synthase *(DXS)* isoforms, a bifunctional squalene synthase (*SQS*) and squalene synthase-like (*SSL*) enzymes working together for the synthesis of the compound. However, its slow growth limits industrial application. In this work, we established the fast-growing diatom *Phaeodactylum tricornutum* as a heterologous chassis for terpenoid production using a synthetic biology Design–Build–Test–Learn (DBTL)-based approach. Episomal uLoop assembly enabled rapid expression and evaluation of *B. braunii DXS* and *SQS* variants, enhancing precursor flux and resulting in the co-accumulation of squalene and carotenoids. This work demonstrates the functional transfer of specialized algal terpenoid enzymes into a tractable diatom host and highlight *P. tricornutum*’s potential as a versatile platform for sustainable, high-value isoprenoid biosynthesis.

## Introduction

Isoprenoids are one of the most diverse and biologically important classes of natural compounds, functioning at the interface of primary and specialized metabolism (Athanasakoglou & Kampranis, 2019; Pruckner et al., 2025; Tetali, 2019). Their structural diversity and bioactivity make them attractive targets in biotechnology, with applications across pharmaceuticals, nutraceuticals, biofuels, and cosmetics. All isoprenoids originate from the polymerization of five-carbon isoprene units.

Carotenoids are tetraterpenoids and give plants, algae, and microorganisms their characteristic yellow to red pigmentation and are widely used in various industrial applications (Esteban et al., 2015; Rodriguez-Concepcion et al., 2018). In humans, they serve as vitamin A precursors, antioxidants, and immunomodulators, contributing to vision and protection against diseases like cancer and cardiovascular disorders (Rodriguez-Concepcion et al., 2018). In photosynthetic organisms, carotenoids are synthesized in the plastid through the methylerythritol phosphate (MEP) pathway, which begins with the rate-limiting enzyme 1-deoxy-D-xylulose-5-phosphate synthase (DXS). Downstream, the condensation of geranylgeranyl diphosphate by phytoene synthase (PSY), followed by a series of desaturation and cyclization reactions, leads to the formation of colored carotenoids with diverse physiological roles (Domonkos et al., 2013; Llorente et al., 2017). Industrial and commercial production through chemical synthesis is costly, energy-intensive, and often yields non-natural isomers (Bogacz-Radomska et al., 2020), while extraction from biomass is limited by land use, seasonality, and waste generation.

Triterpenoids, 30 carbon isoprenoid molecules formed from six isoprene units, include industrially relevant compounds such as sterols and squalene (Sawai & Saito, 2011; Volkman, 2005). Squalene is the universal precursor in sterol biosynthesis and is valued for its use as an emollient, antioxidant, and vaccine adjuvant (Kim & Karadeniz, 2012). Currently, sourcing of squalene for commercial purpose, is based on its extraction from deep-sea shark liver oil, including endangered species, raising environmental and ethical concerns due to overfishing and the ecological vulnerability of shark populations (Dulvy et al., 2014; Popa et al., 2015).

Microalgae offer a promising solution, as being able to naturally synthesize a wide range of carotenoids while certain species and strains naturally accumulate squalene (Lozano-Grande et al., 2018), although in an undomesticated and unoptimized manner, not yet suitable for industrial exploitation. The races B and L of the colonial green microalga *Botryococcus braunii* (Trebouxiophyceae), produces exceptionally high levels of triterpenoids (Boland et al., 2024; Metzger et al., 1985). Races B and L display a metabolism specialized for triterpenoid hydrocarbon synthesis, supported by unique MEP pathway configurations. In race B, genomic and transcriptomic analyses have identified three DXS isoforms: *BbDXS1*, *BbDXS2*, and *BbDXS3* (Matsushima et al., 2012). *BbDXS1* is constitutively expressed and likely maintains plastidial isoprenoid synthesis (e.g., chlorophyll tails, carotenoids), while *BbDXS2* and *BbDXS3* are inducible and strongly upregulated under hydrocarbon-accumulating conditions such as nitrogen limitation or late exponential growth (Matsushima et al., 2012).

In race B, botryococcenes, *B. braunii*’s specific triterpenoids, are produced via a modified squalene biosynthesis pathway: a bifunctional squalene synthase (SQS) catalyzes both the condensation of two FPP molecules into presqualene diphosphate and its subsequent reduction to squalene, while three squalene synthase-like enzymes (SSL-1, SSL-2, SSL-3) partition FPP toward squalene or botryococcene, highlighting metabolic flexibility (Bell et al., 2014; Hirano et al., 2019; Niehaus et al., 2011). By contrast, race L relies on a single SQS enzyme for squalene synthesis (Thapa et al., 2016). Despite its unique metabolic machinery, *B. braunii* grows extremely slowly, forming large mucilaginous colonies with productivities of only 5–10 mg L⁻¹ day⁻¹, limiting its viability for large-scale squalene production (Banerjee et al., 2002; Dayananda et al., n.d.; Nazloo et al., 2024; Wijihastuti et al., 2017). In contrast, the fast-growing and genetically tractable diatom *Phaeodactylum tricornutum* (doubling time ∼12–18 h) has emerged as a promising alternative for squalene biosynthesis and is also a recognized producer of the high-value carotenoid fucoxanthin, highlighting its suitability as a platform for carotenoid accumulation (Kim & Karadeniz, 2012; Xie et al., 2021).

In this study, we applied a systematic synthetic biology approach grounded in the rapid iteration of Design–Build–Test–Learn (DBTL) cycles to develop *P. tricornutum* into a versatile chassis for enhanced terpenoid production. This involved the rapid, modular assembly of episomal constructs via uLoop (Pollak et al., 2019), integration and functional assessment of biosynthetic genes from *B. braunii*, focusing on both upstream precursors and downstream pathways. Differently from approaches relying on random integration, use of episomal vectors in diatoms allowed the rapid generation of complex combinations of genetic parts, coordinated expression and consistent phenotypes. Driven by the hypothesis that *B. braunii’s* DXS and SQS enzymes might be naturally more efficient we focused on specialized BbDXS isoforms, which in *B. braunii* are associated with high flux, specialized terpenoid metabolism (Matsushima et al., 2012), and have the potential to overcome native bottlenecks in precursor supply. By combining these with SQS enzyme variants, we sought to enhance the co-accumulation of both squalene and carotenoids.

Through the iteration of DBTL cycles, we systematically profiled enzyme isoform performance in *P. tricornutum*, host phenotype, and metabolic output, enabling the rapid generation of novel genetic combinations and their phenotypic evaluation. This iterative strategy not only helps understand the functionality of *B. braunii* genes in a heterologous, fast-growing algal host but also provides the demonstration of the suitability of *P. tricornutum* to undergo rapid, iterative DBTL cycles, as a robust and versatile platform for the co-production of high value terpenoids.

## Results and discussion

### The overexpression of *BbDXS* isoforms in *P. tricornutum* results in partial chloroplastic subcellular localization and enhanced pigment amount

To enhance both carotenoid and squalene production, we targeted the upstream segment of the isoprenoid biosynthetic pathway. In *B. braunii* (race B), triterpenoid accumulation relies on the coordinated expression of three DXS isoforms that modulate metabolic flux in response to environmental cues (Matsushima et al., 2012). In this alga, precursors for sterol biosynthesis are supplied by the plastidial MEP pathway, where DXS acts as the first rate-limiting enzyme (Pruckner et al., 2025) and exported into the cytosol and/or ER (Figure 1a).

**Figure 1.**
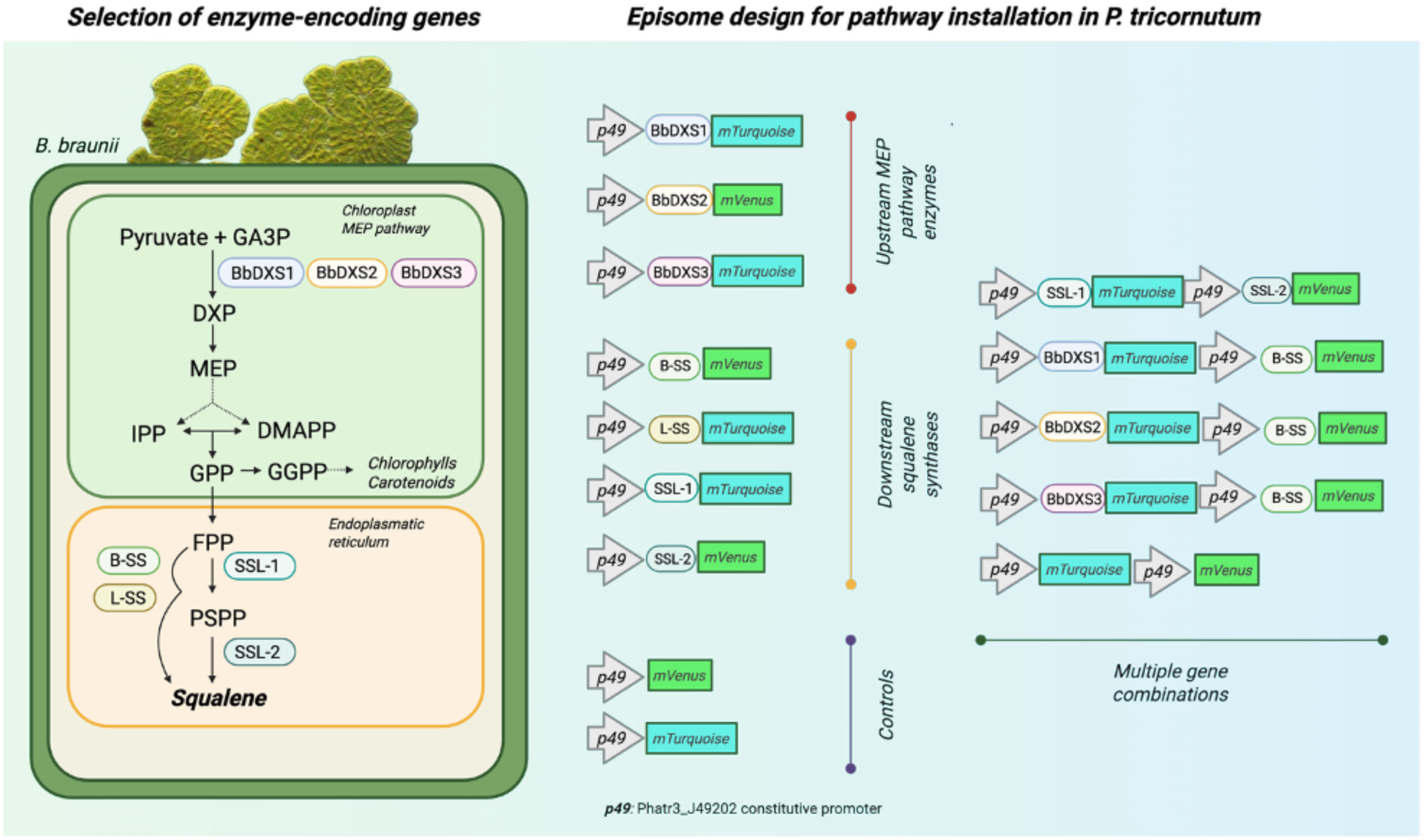
Terpenoid biosynthesis in *B. braunii* and modular expression design in *P. tricornutum*. (A) In *B. braunii*’s chloroplast, the MEP pathway provides IPP and DMAPP, precursors for GPP, GGPP, and FPP. In the ER, FPP is converted to squalene via bifunctional squalene synthases (B-SS, L-SS) or sequentially by SSL-1 and SSL-2 (depending on *B. braunii*’s race). Upstream control involves three DXS isoforms (BbDXS1–3). Dashed lines represent multi-steps reactions. (B) uLoop-based episomal constructs expressed under the constitutive p49 promoter (*Phatr3_J49202* promoter region) fused with fluorescent proteins mTurquoise (mT) or mVenus (mV) at the C-terminal. Single and combinatorial assemblies of upstream and downstream genes enabled functional testing of *B. braunii* enzymes in *P. tricornutum*.

Some evidence indicates that in organisms harboring both the MEP and MVA pathways, intermediates such as IPP and DMAPP can be exchanged between compartments, allowing compensatory adjustments of metabolic flux (Loreto et al., 2004). Based on this, we hypothesized that overexpressing *B. braunii* DXS isoforms in *P. tricornutum* could enhance the plastidial prenyl phosphate pool and thereby increase carotenoid biosynthesis. Similar strategies employing heterologous *DXS* genes, in fact, have successfully increased carotenoid levels in biofortification applications in plants and microalgae (Cen et al., 2022; Estévez et al., 2001). Additionally, the expression of heterologous DXS might circumvent native regulatory bottlenecks in the early steps of isoprenoid biosynthesis, indirectly promoting precursor exchange between the plastidial and cytosolic pools. Although hypothetical, such a mechanism could facilitate metabolic compensation and potentially lead to increased squalene accumulation. *In silico* structural models of PtDXS and BbDXS1–3 generated with AlphaFold2 reveal a highly conserved three-domain architecture typical of DXSs (Fig. S1), consistent with prior biochemical studies (Carretero-Paulet et al., 2006; Matsushima et al., 2012). All four enzymes retain conserved catalytic motifs necessary for thiamine pyrophosphate (TPP)-dependent condensation of pyruvate and glyceraldehyde-3-phosphate, the initiating step of the MEP pathway (Rodriguez-Concepcion et al., 2018). These enzymes show no major structural differences in their catalytic cores, suggesting that if properly localized, could all be functional in *P. tricornutum* (Fig. S1)

Despite their similar enzymatic domains, differences in N-terminal targeting sequences correlated with distinct localization outcomes in *P. tricornutum*. PtDXS features a bipartite signal composed of a ∼25 amino acid ER-type signal peptide (SP) followed by a ∼30–35 amino acid transit peptide (TP) (Fig. S2), consistent with the import requirements of secondary plastids surrounded by four membranes (Felsner et al., 2010; Stork et al., 2013). Plastid import in *P. tricornutum* further relies on a high net positive charge and the presence of key basic residues, typically lysines and arginines, within the TP region (Felsner et al., 2010). In contrast, the TPs of BbDXS1–3, typical of green algae, are longer (∼50–55 amino acids) but lack the SP and often show suboptimal charge distributions, making them poorly suited for diatom plastid targeting (Gruber et al., 2007).

Consistent with these predictions, PtDXS:mVenus localized reliably to the plastid stroma in *P. tricornutum*, while BbDXS1–3:mVenus fusions displayed heterogeneous patterns. BbDXS1 and BbDXS2, in fact, shows a clear structured, periplastidial fluorescence, suggestive of ER surrounding the chloroplast, or outer envelope retention (Liu et al., 2016) to diffuse stromal signal indicative of partial plastid import (Figure 2). This variability reflects the complex import requirements of secondary plastids and the marginal compatibility of *B. braunii* TPs with the diatom’s import machinery (Stork et al., 2013). Notably, even minor variations in TP sequence can affect import efficiency, and the addition of lysine residues has been shown to restore plastid import in heterologous systems (Gruber et al., 2007).

**Figure 2.**
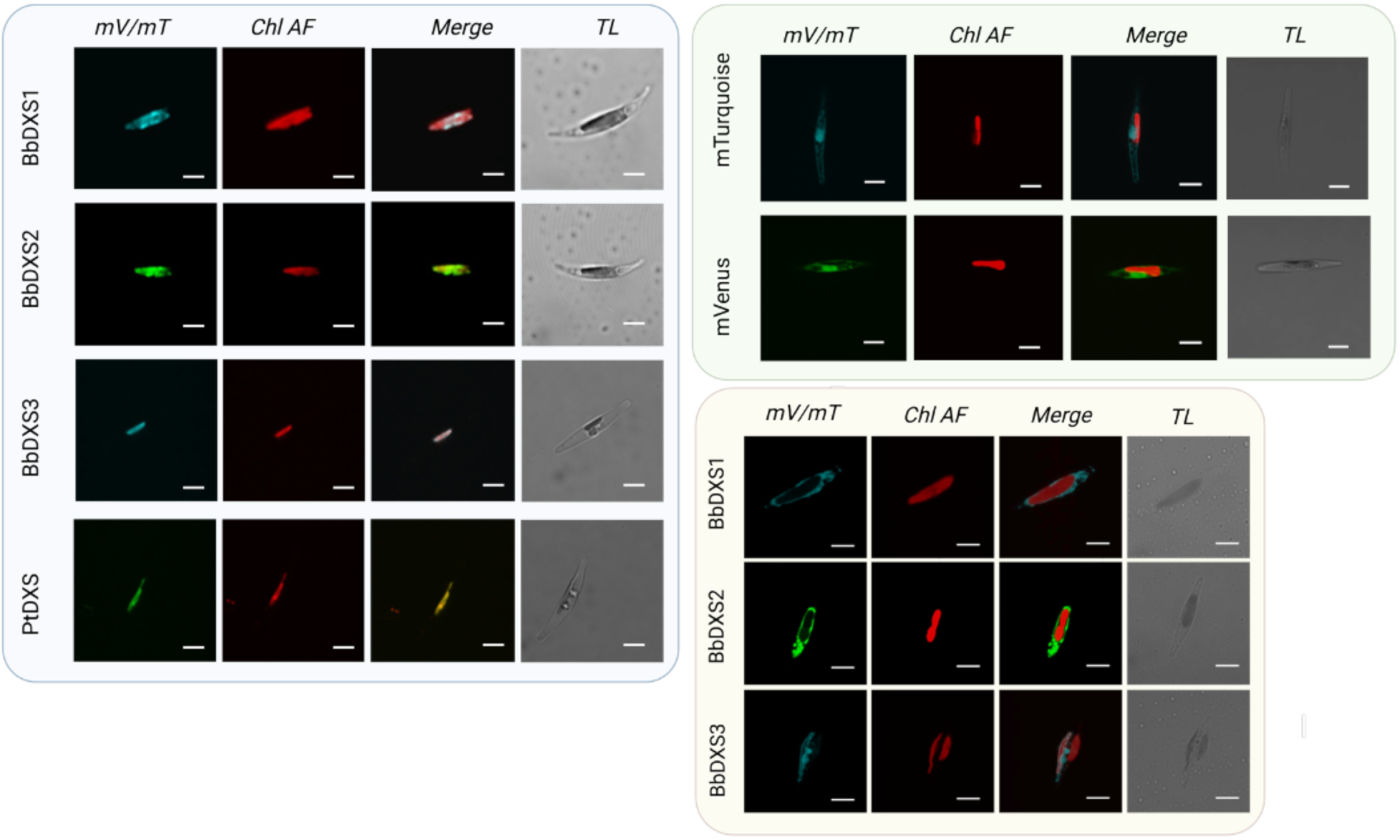
Subcellular localization of *B. braunii* DXS isoforms and PtDXS in *P. tricornutum*. Laser confocal microscopy images show fluorescence from mTurquoise (mT, cyan) or mVenus (mV, green) fused with BbDXS1–3 and PtDXS. Chlorophyll autofluorescence (Chl AF, red) indicates plastid localization. Merged and transmitted light (TL) images illustrate protein distribution relative to plastids and the original shape of the algal cell. White scale bars correspond to 5 µm.

AlphaFold2 models also revealed unique structural features in the mature domain of PtDXS not found in the BbDXS isoforms, including amphipathic helices and hydrophobic patches near the TP cleavage site. These may function as membrane anchors or “stop-transfer” signals, reinforcing PtDXS association with plastid envelope membranes and potentially enhancing import (Felsner et al., 2010). In contrast, BbDXS1–3 appear fully soluble once imported, lacking similar motifs.

Favorable electrostatic conformations or “self-targeting” signals within the mature domain, which typically correspond to positively charged surface residues, may transiently compensate for suboptimal TPs and allow import through the diatom’s plastid translocons (Felsner et al., 2010; Stork et al., 2013) allowing a small but sufficient proportion of BbDXS proteins to reach the stroma and contributing to DXP synthesis.

Despite inefficient targeting, the overexpression of individual *B. braunii DXS* isoforms in *P. tricornutum* showed consistent expression levels (Fig. 3C) and resulted in a consistent increase in chlorophyll *a* (0,401, 0.441 and 0.464 μg/mg DW in BbDXS1, BbDXS2 and BbDXS3, respectively) and carotenoids such as fucoxanthin (1.931, 1.871, 1.917 μg/mg DW) and diadinoxanthin (0.603, 0.642, 0.580 μg/mg DW). These amounts were two-fold higher the ones registered in the mV control and in line with what obtained when overexpressing the endogenous *PtDXS*. (Figure 3A). Similar effects have already been observed in plant and algal systems upon *DXS* overexpression, including a reported 2.4-fold increase in fucoxanthin and a more than 4-fold increase in diadinoxanthin in *P. tricornutum* lines expressing endogenous DXS (Carretero-Paulet et al., 2006; Eilers et al., 2016; Kadono et al., 2015; Manfellotto et al., 2020). This enhancement likely reflects the flux-controlling role of the endogenous PtDXS in the MEP pathway and in the case of BbDXS isoforms overexpression, it is plausible that even partial plastid import can alleviate precursor limitations and drive increased terpenoid biosynthesis (Nisar et al., 2015; Rodriguez-Concepcion et al., 2018)

**Figure 3.**
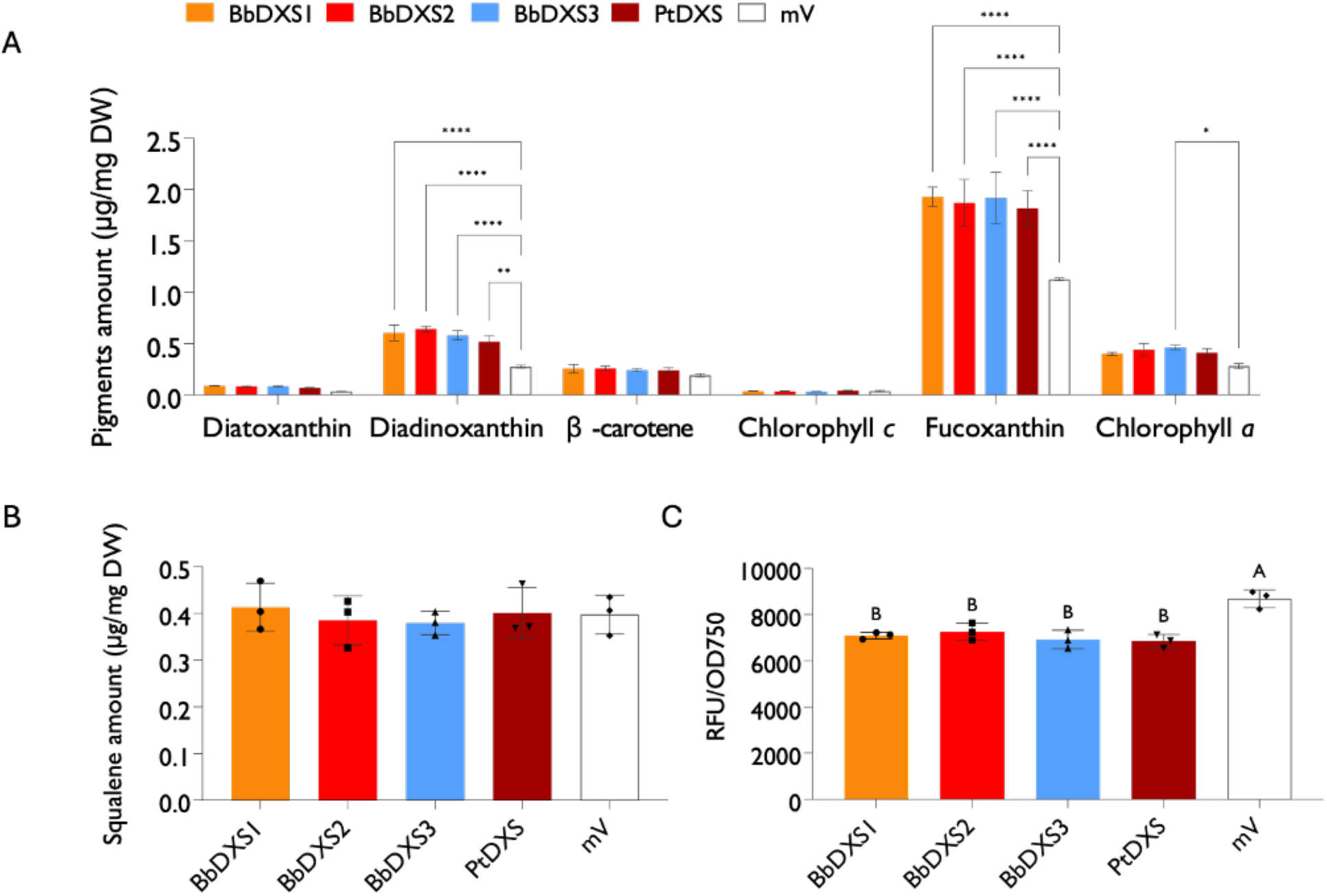
Quantification of pigment and squalene content in *P. tricornutum* lines overexpressing *B. braunii DXS* isoforms. Accumulation of single pigments (A) and squalene (B) in *P. tricornutum* lines overexpressing *B. braunii* different DXS isoforms. Metabolite amounts are expressed in μg/mg of dry weight and calculated with calibration curves of authentic standards. (C) Fluorescence of the respective mVenus or mTurquoise fluorescent protein, used as a proxy for recombinant enzyme abundance at harvest and normalized to optical density. Data represent mean ± SD of three independent biological replicates (n = 3). Statistical significance was assessed using one-way ANOVA followed by Tukey’s test (p < 0.05). Different letters and asterisks indicate statistically significant differences; their absence indicates no significant difference.

Importantly, the expression of BbDXS1–3 did not impair host physiology. Transgenic *P. tricornutum* lines cultivated in presence of Zeocin, showed no detectable growth defects or significant differences in photosynthetic performance compared to control strains after 7 days of culture (Figure 4A, 3B), indicating that the metabolic load of heterologous DXS expression is well tolerated. The diatom cell lines accumulating higher amounts of diadinoxanthin, also showed a measurable enhancement in non-photochemical quenching (NPQ) under high light stress associated then with a faster recovery of the built up NPQ once the cultures were re-exposed to darkness (Figure 4C, D), compared to the control, suggesting that the expanded xanthophyll cycle pigment pool (i.e., specifically more diadinoxanthin available for conversion into diatoxanthin) contributes to improved photoprotection. NPQ is a photoprotective mechanism that dissipates excess absorbed light energy as heat, preventing photodamage to the photosynthetic machinery. In diatoms, NPQ is closely linked to the diadinoxanthin (Dd)–diatoxanthin (Dt) cycle, where light-induced de-epoxidation of Dd to Dt enhances NPQ efficiency by facilitating energy dissipation in the thylakoid membranes. The reversible conversion between these pigments allows dynamic regulation of NPQ in response to changing light conditions (Goss et al., 2006; Grouneva et al., 2009; Kagatani et al., 2022; Lavaud et al., 2007).

**Figure 4.**
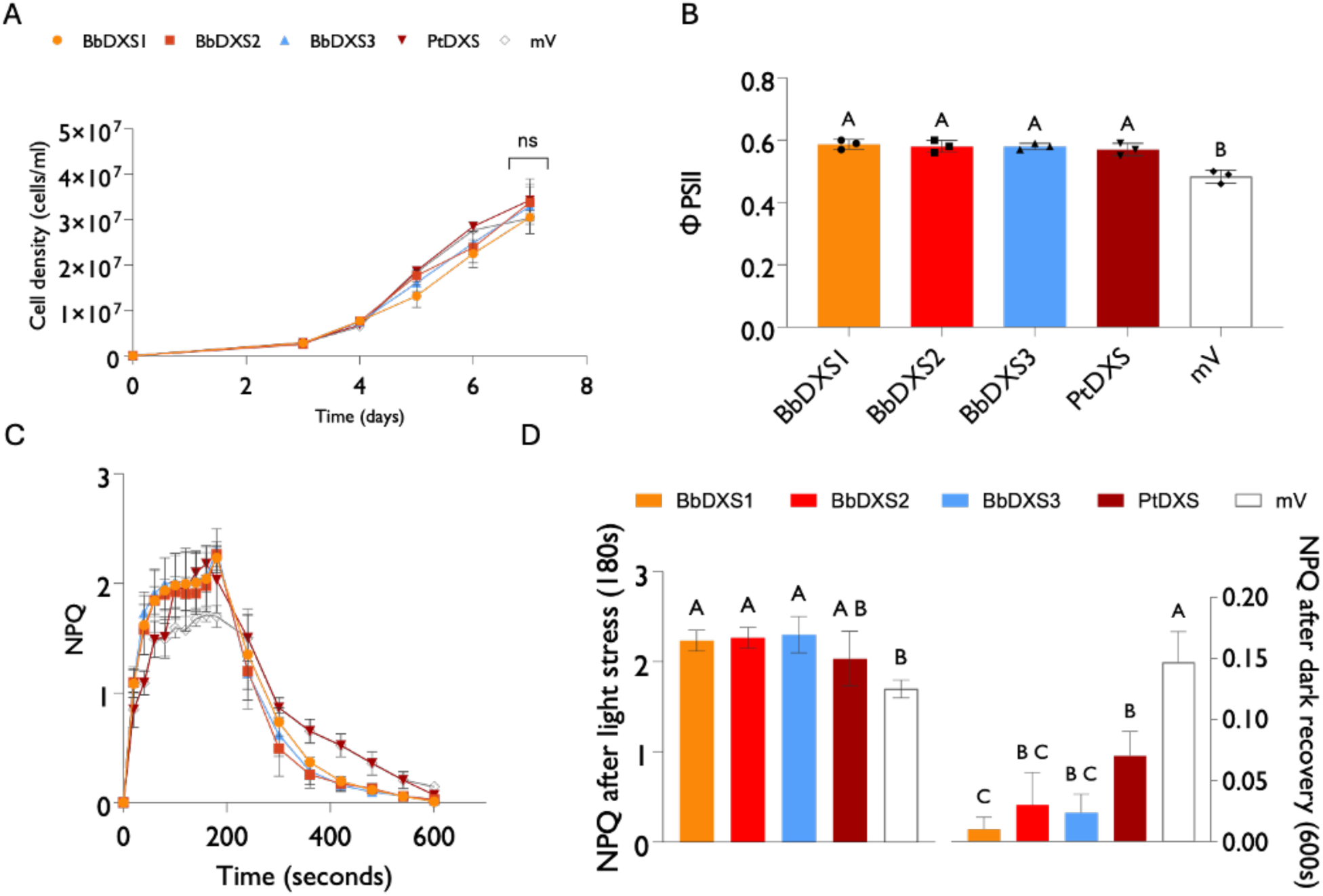
Growth and photosynthetic performance of *P. tricornutum* lines expressing *B. braunii* DXS isoforms. (A) Growth curves of transgenic lines overexpressing *BbDXS1–3* and *PtDXS,* expressed as cell density calculated through flow cytometry (B) Effective quantum yield (ϕPSII) in the same lines (C) NPQ (non-photochemical quenching) induction and relaxation kinetics under high light (HL, 600 µmol m⁻² s⁻¹) followed by darkness. (D) Bar plots showing the NPQ values observed after 200 s in high light and after 600 s in recovery, respectively. Data represent mean ± SD from three independent biological replicates (n = 3). Different letters indicate statistically significant differences (one-way ANOVA, *p*<0.05).

While earlier studies did not report NPQ changes in DXS-overexpressing lines (Kadono et al., 2015) our results link the effect of the overexpression of increased xanthophyll cycle pigment pools to enhanced NPQ capacity. Notably, the amount of chlorophyll remained unchanged, indicating that excess carotenoids were likely stored outside pigment-protein complexes, possibly in lipid bodies (Eilers et al., 2016). This supports the notion that overexpressing heterologous *DXS*, despite the complex regulation which somehow limits the maximum yield of carotenoids reachable, can effectively shift metabolic flux toward photoprotective carotenoids, reinforcing light stress resistance without compromising photosynthetic core components and resulting in an algal strain with a potential improved stress response.

Squalene levels in *P. tricornutum* cell lines overexpressing either isoform of DXS remained unaltered compared to the control (0.413, 0.384, 0.379, 0.400 μg/mg DW for BbDXS1, BbDXS2, BbDXS3 and PtDXS respectively, compared to the 0.397 μg/mg DW found in the mV control), most likely due to the plastidial compartmentalization of the MEP pathway and carotenoid biosynthesis in *P. tricornutum*, with squalene and sterol biosynthesis primarily relying on precursors from the MVA pathway (Jaramillo-Madrid, Abbriano, et al., 2020; Jaramillo-Madrid, Ashworth, et al., 2020).

### *B. braunii* SQS’s are compatible with the machinery of *P. tricornutum* and efficiently increase the amount of squalene and downstream sterols

To enhance squalene production in *P. tricornutum*, we individually overexpressed four *B. braunii* genes directly involved in triterpene biosynthesis: squalene synthase from race B (*B-SS*), squalene synthase from race L (*L-SS*), and two squalene synthase-like genes from race B (*SSL-1* and *SSL-2*). We hypothesized they might bypass diatom regulatory constraints and operate more efficiently, given *B. braunii*’s high squalene productivity. AlphaFold structural models show that both B-SS and L-SS display a compact, globular architecture with a well-defined C-terminal transmembrane helix, consistent with predicted membrane anchoring and localization to the ER, as expected for canonical squalene synthases (Figs. S3, S4) (Niehaus et al., 2011; Okada et al., 2004). In contrast, SSL-1 lacks such clear membrane-spanning domains and exhibits more extended, modular topologies with surface-exposed regions potentially involved in protein–protein interactions, supporting previous biochemical findings that SSL-1 does not act independently but requires partner proteins like SSL-2 or SSL-3 for full catalytic activity (Niehaus et al., 2011). The observed structural traits are consistent with a multi-enzyme complex model, where proper spatial organization and cofactor interactions are necessary for metabolite channeling and function. Laser confocal microscopy analyses showed that B-SS, L-SS, SSL-1, and SSL-2 predominantly localized to the ER likely for the presence of a transmembrane domain in these proteins (Niehaus et al., 2012) (Fig. 5).

**Figure 5.**
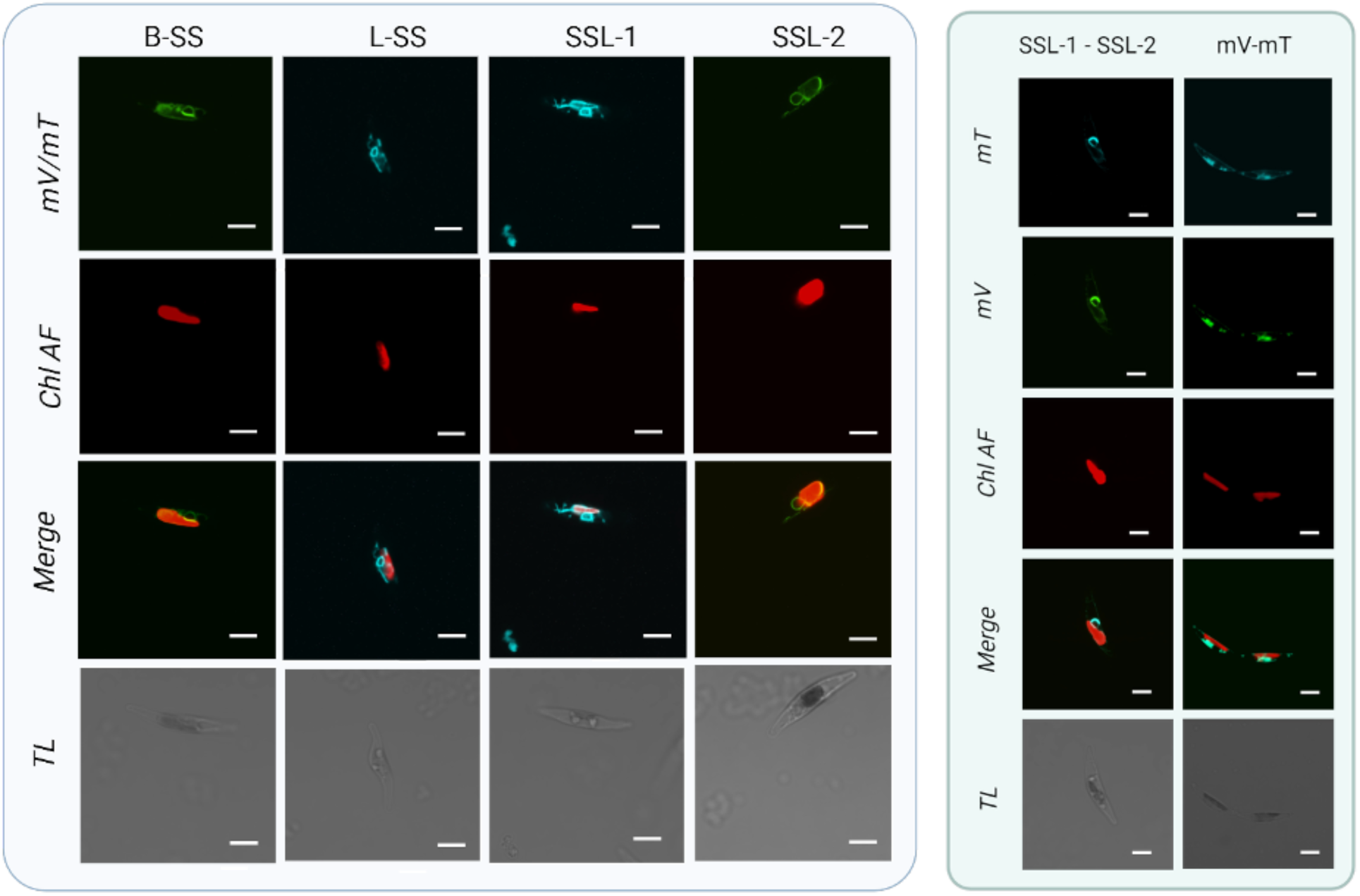
Subcellular localization of *B. braunii* B-SS, L-SS, SSL-1 and SSL-2 in *P. tricornutum*. Laser confocal microscopy images show fluorescence from mTurquoise (cyan, mT) or mVenus (green, mV) fused with the different proteins. Chlorophyll autofluorescence (Chl AF, red) indicates plastid localization. Merged and transmitted light (TL) images illustrate protein distribution relative to plastids and the original shape of the algal cell. White scale bars correspond to 5 µm.

In eukaryotes, sterol biosynthesis occurs in the ER, and this is likely also the case in *P. tricornutum* (Jaramillo-Madrid et al., 2020; Volkman, 2005). The precise localization of squalene biosynthesis in this organism remains unresolved as the PtIDISQS fusion enzyme (isopentenyl diphosphate isomerase–squalene synthase) lacks a clear signal peptide or transit sequence (Fabris et al., 2014), leaving its subcellular targeting ambiguous. However, the alternative squalene epoxidase (AltSQE) functions in the ER (Pollier et al., 2019), suggesting that, at least squalene conversion takes place in this compartment.

Overexpressing *B-SS* and *L-SS* resulted in significantly higher squalene amounts compared to the mV control (0.788 and 0.717 μg/mg DW, corresponding, for our set-up to 3.47 mg/L and 3,15 mg/L, against 1.68 mg/L) with *B-SS* being the most effective (Fig. 6A). In contrast, expression of *SSL-1* or *SSL-2* alone had no significant effect. SSL-1 catalyzes the first step of squalene biosynthesis, forming presqualene diphosphate (PSPP) from two FPP molecules, but requires SSL-2 to complete the reaction. Likewise, SSL-2 functions only in conjunction with SSL-1 and has not metabolic function on its own (Niehaus et al., 2011). Accordingly, the co-expression of SSL1 and SSL2 in *P. tricornutum* restore full enzymatic functionality with levels of squalene accumulation like the ones obtained when expressing *B-SS* (0.776 μg/mg DW corresponding to 3,41 mg/L) (Fig. 6).

**Figure 6.**
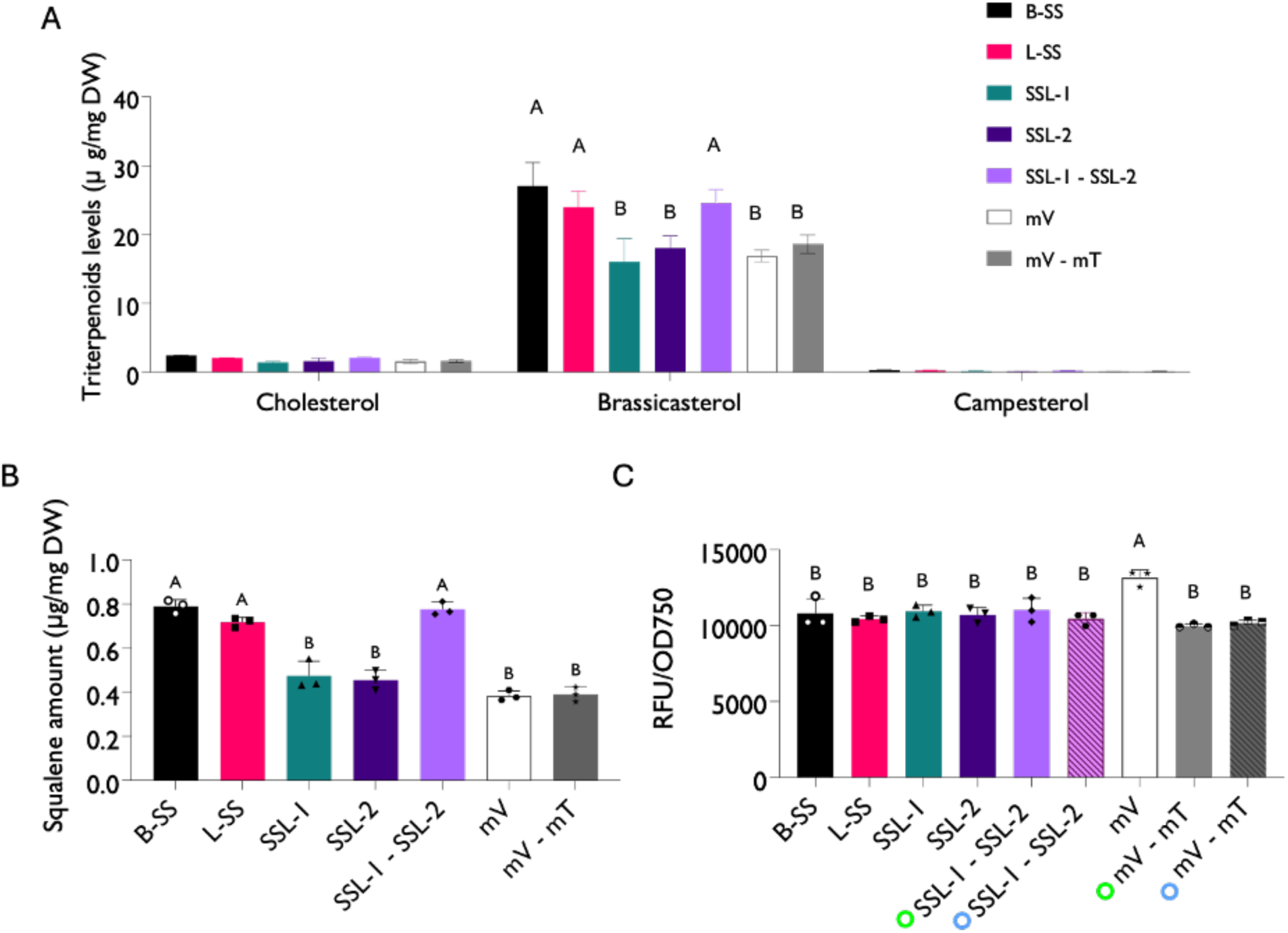
- Quantification of triterpenoid content in *P. tricornutum* lines overexpressing expressing *B. braunii* different squalene synthases. Levels of the single triterpenoids (A) and squalene (B) in *P. tricornutum* cell lines overexpressing *B. braunii* different SQS. Metabolite levels are expressed as μg/mg of dry weight and calculated with calibration curves of authentic standards. (C) Fluorescence of the respective mVenus or mTurquoise fluorescent protein (indicated by the coloured circles), used as a proxy for protein production and normalized to optical density. Data represent mean ± SD of three independent biological replicates (n = 3). Statistical significance was assessed using one-way ANOVA followed by Tukey’s test (p < 0.05). Different letters and asterisks indicate statistically significant differences; their absence indicates no significant difference. (

Although overall squalene levels in the engineered lines remained low, their increase relative to controls provides valuable information for guiding further optimization within the DBTL cycle iterations. Increasing sterol content (Fig. 6B) indicates downstream conversion of squalene. Unlike previous studies in *P. tricornutum*, where overexpression of *tHMGR* or *N. oceanica SQE* caused transient squalene accumulation without affecting total sterol levels (Jaramillo-Madrid et al., 2020), the use of *B. braunii* isoforms here additionally supports the flux toward sterol synthesis resulting in increased brassicasterol amounts in the B-SS, L-SS and SSL-1 – SSL-2, overexpressing lines, likely produced from the additional squalene.

Interestingly, diatoms overexpressing all *BbSQS* versions grew less compared to the respective control and showed a lower photosynthetic activity with a reduction negatively correlated to the respective increase of squalene and sterols (Figure 7A, 7B). Sterols, especially cholesterol are involved in membrane stability and fluidity (Volkman, 2005, 2016) and shifts in sterol composition (e.g., higher phytosterols and lower cholesterol) can reduce photosynthetic efficiency by altering thylakoidal membrane restructuring and impacting transient photosynthetic performance (H. Schaller, 2003; S. Schaller et al., 2011). As an additional potential link between altered sterol profiles and destabilization of diatom photophysiology, we observed that, unlike the DXS-overexpressing lines, the SQS lines exhibited NPQ levels similar to the control under high light (Figure 7C, D), followed by a delayed recovery upon return to darkness, likely reflecting a slower reorganization of internal thylakoid membranes (Grosjean et al., 2015)

**Figure 7.**
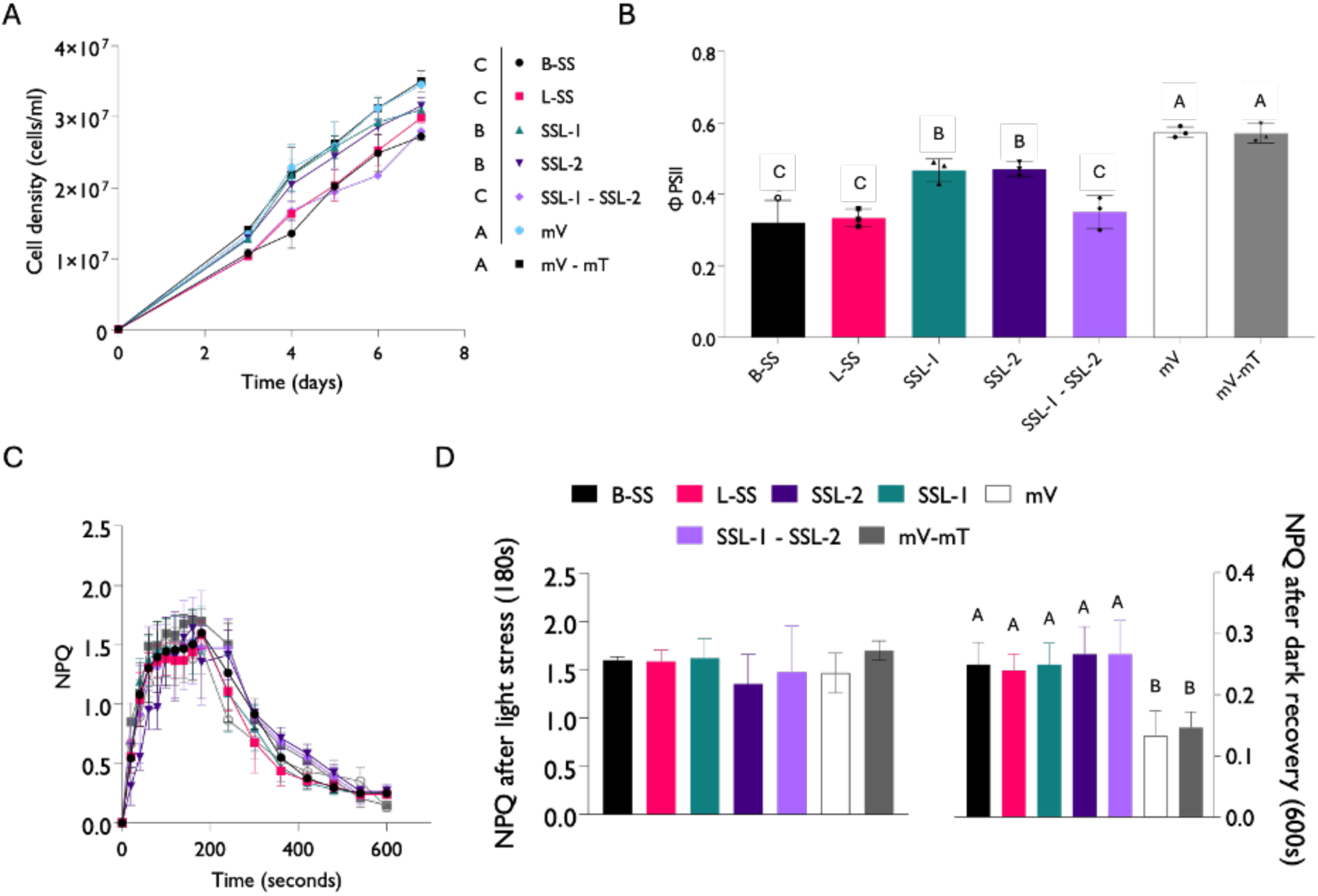
Growth and photosynthetic performance of *P. tricornutum* lines expressing various configurations of *B. braunii SQS*. (A) Growth curves of transgenic lines overexpressing *B-SS, L-SS, SSL-1, SSL-2 and SSL-1 – SSL-2*, expressed as cell density calculated through flow cytometry (B) Effective quantum yield (ϕPSII) in the same cell lines (C) NPQ (non-photochemical quenching) induction and relaxation kinetics under high light (HL, 600 µmol m⁻² s⁻¹) followed by darkness. (D) Bar plots showing the NPQ values observed after 200 s in high light and after 600 s in recovery, respectively. Data represent mean ± SD from three independent biological replicates (n = 3). Different letters and asterisks indicate statistically significant differences; their absence indicates no significant difference. (one-way ANOVA, *P*<0.05).

### Intracellular lipid droplets increase in number with SQS overexpression but cannot be used as an efficient storage structure for squalene

SQS-overexpressing *P. tricornutum* lines showed increased sterol content and increased lipid droplets (LD) accumulation (Figure 8A), detected via CARS microscopy targeting the ∼2845 cm⁻¹ CH₂ stretch of neutral lipids. In *Saccharomyces cerevisiae*, genetic manipulation of ER-located enzymes can lead to drastic changes in this organelle morphology expanding it over its natural conformation (Arendt et al., 2017). We thus hypothesize that the abundance of SQS in ER membranes, or the elevated sterol levels, may contribute to membrane destabilization within the ER. Although this may not alter ER morphology, it could promote the formation of other lipid-based structures such as lipid droplets LD. The formation of these structures is commonly associated with metabolic stress or nutrient starvation, conditions that lead to triacylglycerol (TAG) accumulation (Guéguen et al., 2021; Maeda et al., 2017; Sayanova et al., 2017). In yeast, lipid droplets derived from the ER serve as storage sites for neutral lipids and isoprenoid intermediates like squalene which accumulation is closely linked to lipid droplets proliferation, reflecting an adaptive response to increased metabolite production (Spanova et al., 2012; Ta et al., 2012).

**Figure 8.**
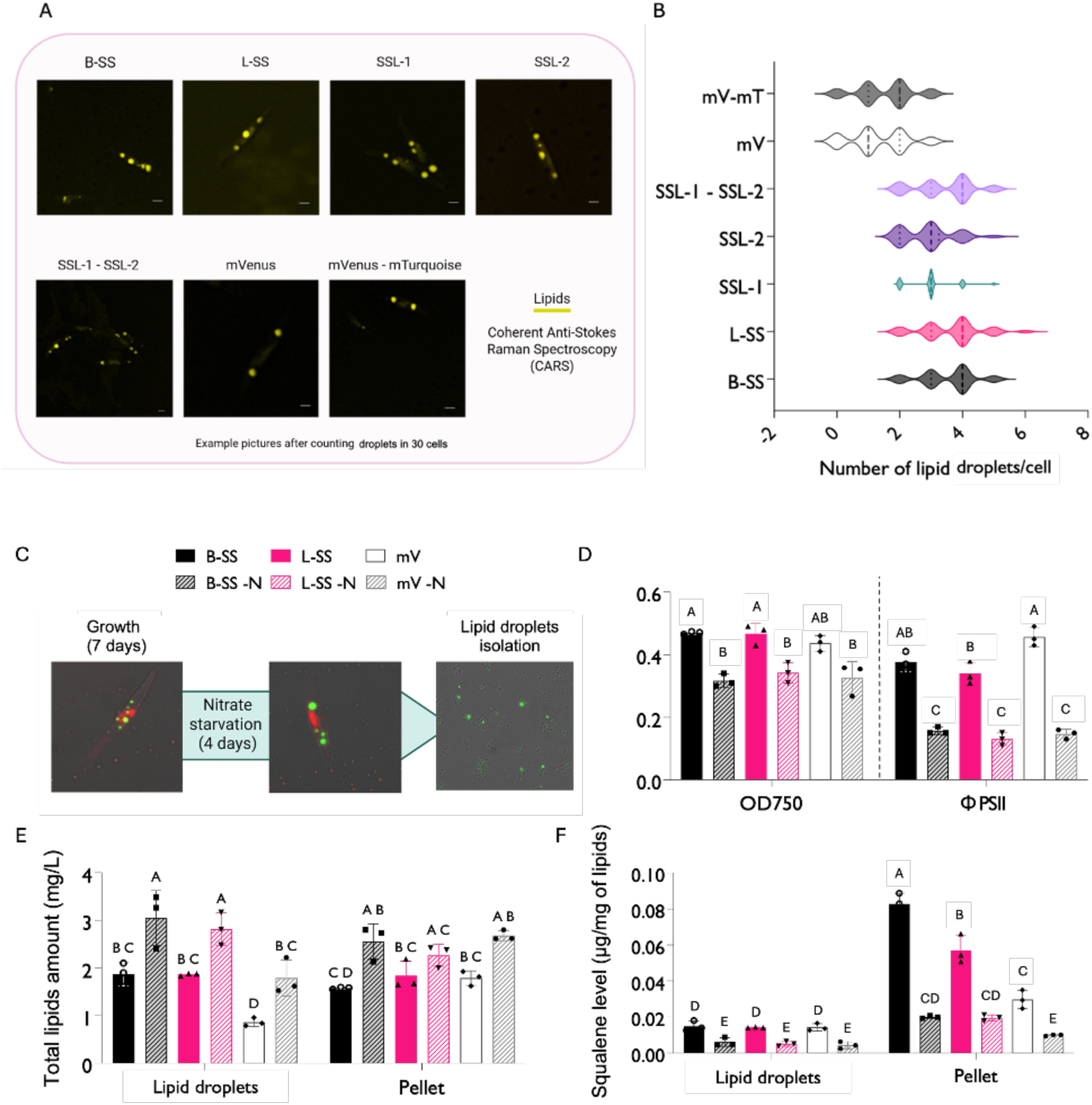
Lipid droplets accumulation and squalene distribution in *P. tricornutum* cell lines expressing *B. braunii* squalene synthase (SQS) and squalene synthase-like (SSL) enzymes. (A) Representative Coherent Anti-Stokes Raman Scattering (CARS) microscopy images show lipid droplets (yellow) in cells expressing B-SS, L-SS, SSL-1, SSL-2, and SSL-1 + SSL-2, compared with fluorescent protein controls (mVenus, mTurquoise, and mVenus–mTurquoise). Scale bars: 2 µm. The violin plot (B) summarizes lipid body counts per cell from 30 individual cells per line. The width of each distribution represents the frequency of observations, with wider sections indicating more common values. Horizontal bars denote medians, and thin lines indicate interquartile ranges. Increased median and broader distributions reflect higher and more heterogeneous lipid accumulation in specific constructs. (C) Schematic illustration of the experimental setup: cells were grown for 7 days under nutrient-replete conditions, followed by 4 days of nitrate starvation before lipid body isolation. Confocal images show chlorophyll autofluorescence (red) and lipids stained with Nile Red (green). (D) Optical density and effective quantum yield used as a phenotypic proxy of the effect of nitrogen starvation on the culture (E) Quantification of total lipids in the isolated lipid droplets and in the remaining pellet (F) amount of squalene found in the same two cellular spaces, relative to the amount of total lipids. Data represent mean ± SD from three independent biological replicates (n = 3). Different letters indicate statistically significant differences (one-way ANOVA, *P*<0.05).

The cell lines that accumulated the highest levels of squalene (B-SS and L-SS) also displayed the greatest number of lipid droplets (Figure 8B). Given this correlation and considering previous reports of small amounts of squalene present in *P. tricornutum* lipid droplets, we investigated whether these could serve as sequestration hubs for squalene and prevent its conversion into downstream sterols, not only when their formation is the result of direct squalene accumulation but also when their formation is triggered by other physiological conditions. A similar approach has been demonstrated in plants, where inducing plastoglobuli (i.e., lipid-rich plastid subcompartments that store carotenoids and other hydrophobic molecules) proliferation through environmental stimuli such as high light supports carotenoid metabolic engineering strategies, likely by increasing plastid storage capacity. (Morelli, Romañach, et al., 2023; Morelli, Torres-Montilla, et al., 2023; Wijk & Kessler, 2017).

To test this hypothesis, we induced LD proliferation via nitrate starvation in the cell lines which showed higher productivity of squalene (Yang et al., 2013). Nile Red staining confirmed an increase in both the size and number of lipid bodies, while reduced photosynthetic activity and growth rate validated the effectiveness of the stress treatment (Fig. 8D). We then quantified total lipid content in both the isolated lipid body fraction and the remaining pellet, containing membranes and cell debris. As expected, nitrate starvation induced an increase in the amount of total lipids in all the cell lines expressing SQSs but, compatibly with the previous microscopy observation, lines overexpressing heterologous SQSs showed increased lipid content in the lipid bodies fraction compared to the respective controls, suggesting enhanced lipid bodies formation (Figure 8E). Although small amounts of squalene have been previously detected in the LDs of *P. tricornutum*, our results indicate that the excess squalene does not accumulate within these structures. Instead, most of the squalene remained in the pellet fraction, suggesting its predominant localization to membrane-associated compartments (Fig. 8F). This observation aligns with earlier lipidomic analyses reporting that *P. tricornutum* LDs are mainly composed of neutral lipids, with minimal squalene content (Leyland et al., 2020; Lupette et al., 2019).

These findings imply that, unlike in other systems where LD proliferation correlates with metabolite buffering, increasing LD size alone in *P. tricornutum* is insufficient to divert squalene from its rapid conversion into sterols or from membrane association because the absence of a native sequestration role for LD may limit their capacity to act as an effective metabolic sink. Consequently, our “push and pull” strategy consisting in combining metabolic enhancement with potential storage in lipid bodies, appears unfeasible in this system. Instead, the observed lipid body increase is more likely a secondary effect of stress responses associated with sterol accumulation and reduced photosynthetic performance.

### The combinatorial overexpression of BbDXS isoforms with B-SS enhances the metabolic flux towards both carotenoids and squalene in *P. tricornutum*

Based on the outcomes of the previous strategies, we designed the next DBTL cycle iteration by combining the overexpression of each *B. braunii DXS* isoform with the best-performing squalene synthase, *B-SS*. The rationale was to synergistically boost both carotenoid and squalene biosynthesis by enhancing precursor availability and potentially improving photosynthetic performance via DXS-mediated fine-tuning of plastidial isoprenoid metabolism. Furthermore, we hypothesized that increasing flux through the upstream MEP pathway could alleviate metabolic stress associated with high squalene synthesis, ultimately supporting a more robust production chassis. Upon co-expression of each *DXS* isoform with *B-SS*, we observed that the distinct subcellular localization patterns previously seen when expressed individually (mostly in the chloroplast for DXS, in the ER for B-SS) were maintained. This indicates that their concurrent expression does not interfere with proper targeting, supporting their compatibility in engineered pathways (Fig. 10A). Furthermore, an increase in growth was observed, accompanied by a partial recovery of the effective quantum yield of photosystem II, reaching levels comparable to those of the control lines. This improvement is likely attributable to the effect of *DXS* overexpression, consistent with the response observed when this gene was expressed individually. (Fig. 10B, C).

**Figure 9.**
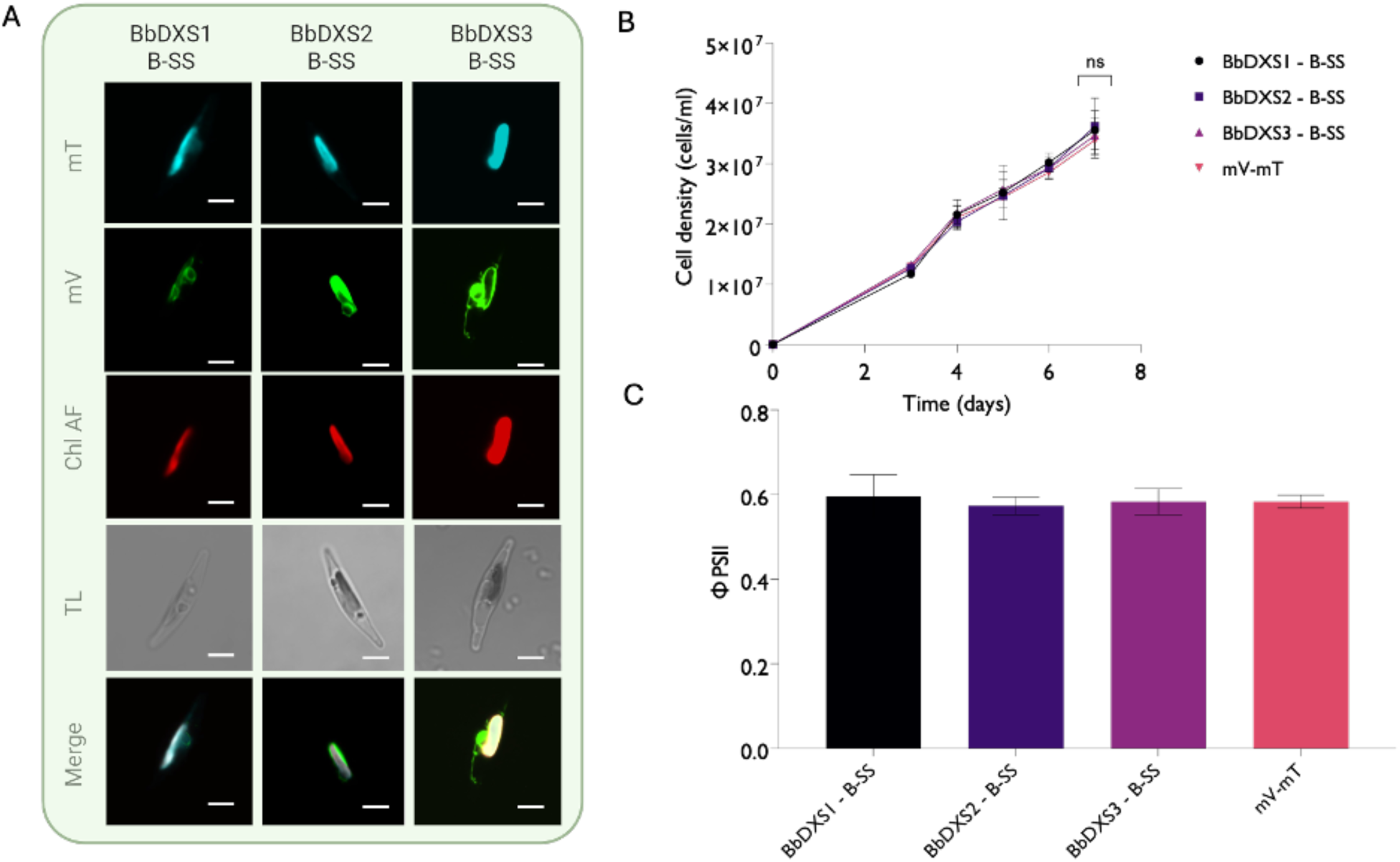
Overexpression of *B. braunii DXS* isoforms in *P. tricornutum* in combination with B-SS. (A) Confocal images show fluorescence from mTurquoise (cyan, mT) or mVenus (green, mV) fused with the different proteins. Chlorophyll autofluorescence (Chl AF, red) indicates plastid localization. Merged and transmitted light (TL) images illustrate protein distribution relative to plastids and the original shape of the algal cell. White scale bars correspond to 5 µm (B) Growth curves of transgenic lines expressed as cell density calculated through flow cytometry (B) Effective quantum yield (ϕPSII) in the same lines. Data represent mean ± SD from three independent biological replicates (n = 3). Different letters and asterisks indicate statistically significant differences; their absence indicates no significant difference (one-way ANOVA, P<0.05).

**Figure 10.**
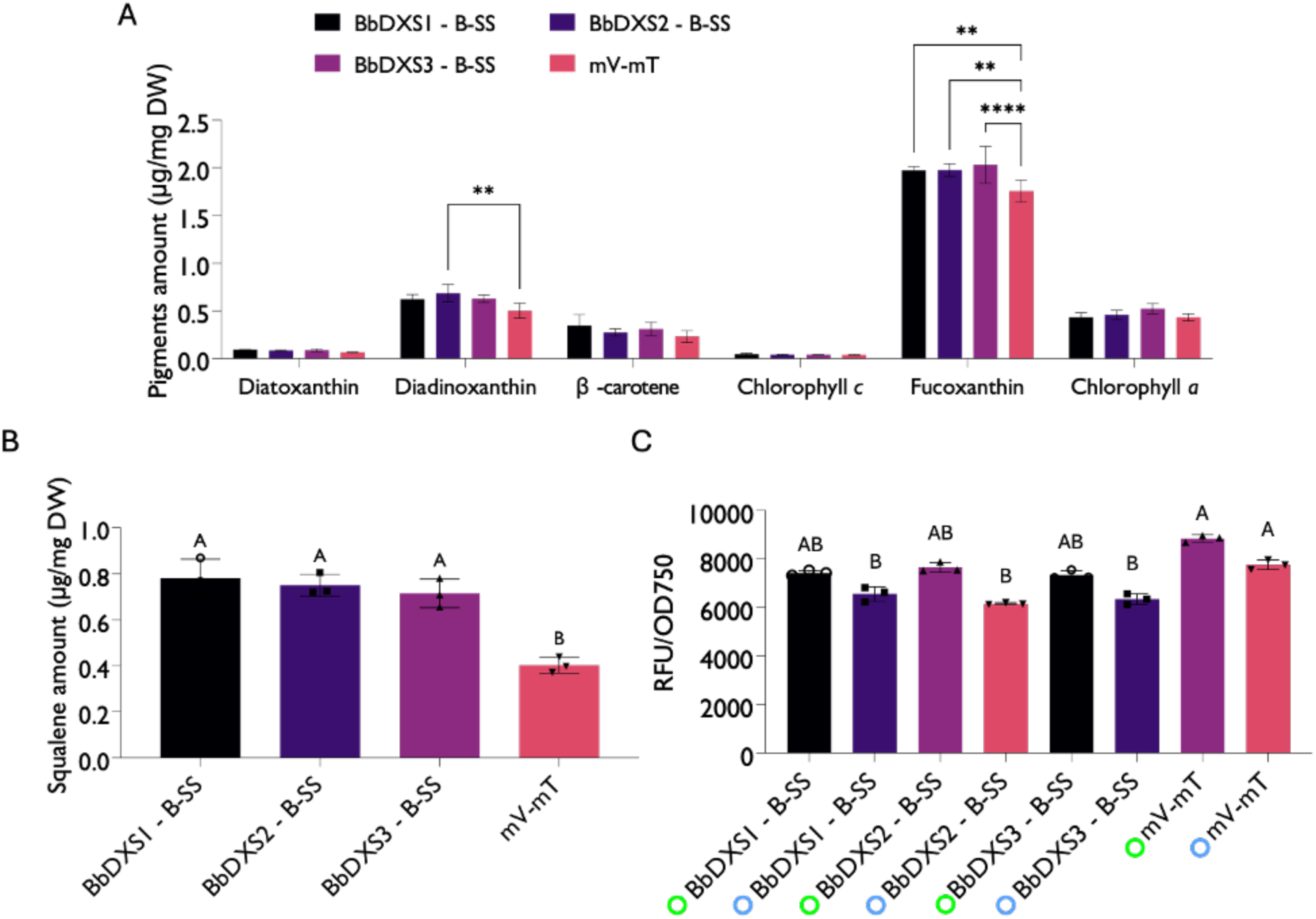
Quantification of pigment and squalene content in *P. tricornutum* cell lines overexpressing *B. braunii DXS* isoforms in combination with *B. braunii’s* B-SS. Levels of the single pigments (A) and squalene (B) in *P. tricornutum* cell lines overexpressing *B. braunii* different DXS isoforms together with B-SS enzyme. Metabolite levels levels are expressed as μg/mg of dry weight and calculated according to a calibration curve of the authentic standards. (C) Fluorescence of the respective mVenus or mTurquoise fluorescent protein (indicated by the coloured circle), used as a proxy for protein production and normalized to optical density. Data represent mean ± SD of three independent biological replicates (n = 3). Statistical significance was assessed using one-way ANOVA followed by Tukey’s test (p < 0.05).

These combinatorially engineered diatom cell lines showed elevated carotenoid content, comparable to those detected in *DXS*-overexpressing lines (Fig. 3). All the three lines accumulates higher amounts of fucoxanthin compared to the mV-mT control (1.97, 1.97 and 2 μg/mg DW for BbDXS1, BbDXS2 and BbDXS3 – B-SS, respectively) compared to the mV-mT control (1.75 μg/mg DW). Squalene levels also increased in the three lines, although not beyond those observed with *B-SS* overexpression alone (0,780, 0.750, 0.714 μg/mg DW, corresponding, for our set-up, to 3.43 mg/L, 3.3 mg/L, and 3.14 mg/L) and resulting in a 1.5-fold increase compared to the amount of squalene found in the mV-mT control (0,402 μg/mg DW, corresponding to 1,76 mg/L) (Figure 11A, 11B).

These results suggest that the combined production of *B. braunii*’s three DXS isoforms and of B-SS has effects that extend beyond simple additive increases in metabolite production. In fact, co-expression may partially mitigate the physiological imbalances or “damage” caused by the overexpression of a single heterologous protein (e.g., the B-SS). Although the combination of DXS and SQS does not produce higher overall carotenoid or squalene yields compared with expressing either gene alone, the observed increase in photoprotective capacity suggests that these hybrid strains may be better adapted to fluctuating or high-light environments, potentially supporting improved long-term productivity.

## Conclusions

This study demonstrates the feasibility and value of a synthetic biology-driven DBTL approach for engineering complex isoprenoid metabolism in *P. tricornutum*. By overexpressing the three *DXS* isoforms and the different *SQS* from *B. braunii*, we gained new insights into both the localization behavior and functional integration of non-native enzymes in a diatom host. Through this process, we identified B-SS from *B. braunii* as promising enzymatic target that effectively allows to increase squalene and downstream sterols levels. Wild-type *P. tricornutum* contains negligible free squalene so the amount obtained by overexpressing *B-SS* and *L-SS* (0.788 and 0.717 µg/mg DW) were on par with or higher than prior examples. In fact, trials of overexpression of key sterol pathway genes (e.g., HMGR) (Jaramillo-Madrid et al., 2020) were shown to boost squalene accumulation by an order of magnitude relative to wild type, but still not exceeding 0.5 μg/mg DW. In comparison to *P. tricornutum*, microbial hosts lacking downstream sterol pathways can accumulate squalene more effectively. *E. coli*, which lacks triterpenoids, can stockpile large amounts of heterologously produced squalene with minimal diversion to other compounds (Gohil et al., 2019) while a *Synechocystis* cyanobacterium mutant (Δ*shc*, deficient in squalene-hopene cyclase) reached 14 mg/L squalene by overexpressing squalene synthase (Germann et al., 2023). By contrast, eukaryotic systems such as yeast and green algae require a higher degree of genetic intervention to avoid squalene conversion to downstream sterols. Engineered *S. cerevisiae* strains, for example, have achieved squalene titers of 0.75 g/L in shake flasks (exceeding 2 g/L in fed-batch fermentations) by overexpressing pathway genes and partially inhibiting squalene epoxidation (Han et al., 2018) whereas in *Chlamydomonas reinhardtii* even a partial knockdown of squalene epoxidase was needed to obtain 1.1 mg/L squalene (Kajikawa et al., 2015). Coherently, and considering that in our work no intervention was made to avoid diversion of squalene towards the downstream pathway, our engineered *P. tricornutum* strains accumulate comparatively little squalene. Even so, the combinatorial overexpression of *B. braunii* DXS isoforms with the high-activity squalene synthase (B-SS) proved beneficial by markedly increasing squalene amount and carotenoid synthesis, thus improving photoprotection.

Despite this, methodologically, this work illustrates the value of an iterative DBTL pipeline for algal engineering. Using episome-based modular vectors together with high-throughput flow-cytometric screening allowed rapid prototyping of transgenic lines with controlled copy number and expression. Extrachromosomal maintenance avoids insertional mutagenesis and variegation, yielding more uniform expression (George et al., 2020). Combined, episomal expression, modular DNA assembly, and high-throughput phenotyping provide a robust platform for accelerated strain development. This enables *P. tricornutum* to be engineered efficiently for enhanced high value compounds biosynthesis, narrowing the performance gap with established yeast and bacterial hosts.

## Materials and methods

### Algae strains and cultivation

Axenic cultures of *P. tricornutum* CCAP1055/1 were obtained from the CCAP (Culture Collection of Algae and Protozoa, SAMS Research Services Ltd., Scottish Marine Institute, Oban, U.K., http://www.ccap.ac.uk) and were grown in liquid Enriched Seawater, Artificial Water (ESAW) medium (Berges et al., 2001). For experiments, diatoms producing recombinant proteins were cultured in individual 50 ml flasks in fully controlled Innova® S44i shaking incubators (Eppendorf) under continuous light regime (100 μE m^-2^sec^-1^) at 21°C, shaking at 95 rpm. Parameters such as optical density, cell count, and fluorescent protein intensity (used as a proxy for recombinant enzyme production) were monitored on days 3, 4, 5, 6, and 7 from the start of the experiment.

For nitrate depletion conditions aimed at increasing lipid droplet size, diatoms were cultured for 7 days under the abovementioned conditions in 250 mL volumes. Cells were then gently centrifuged at 3500 g for 10 minutes and resuspended in an equal volume of ESAW medium lacking nitrate. Cultures were maintained under nitrogen-depleted conditions for 4 days prior to further processing. Control algae were resuspended in unmodified ESAW medium. A measure of optical density and effective quantum yield was used to confirm the effect of the nitrate depletion. Intracellular lipid bodies were stained and visualized by Nile red staining (see “Lipid bodies isolation section”)

### Cloning, constructs assembly and diatom transformation by bacterial conjugation

All episomes used in this study were assembled using the *u*Loop assembly method (Pollak et al., 2019). Individual components for episome assembly (L0 parts) were constructed and domesticated following the *u*Loop assembly syntax (Table S1). Assembly reactions were carried out using the corresponding *u*Loop backbone vectors for each assembly level, as described by Pollak et al. Following domestication, each L0 part was verified by Sanger sequencing. DNA parts used for domestication were codon-optimized, designed with the appropriate overhangs, synthesized de novo, and purchased from Genewiz (Azenta Life Sciences). The complete sequences of all L0 plasmids in Table S2. Expression cassettes included genes involved in the squalene biosynthetic pathway (*B-SS, L-SS, SSL1, SSL2* annotated as *AF205791*, *KT388100*, *HQ585058* and *G0Y287*, respectively), the three isoforms of *B. braunii* DXS (*BbDXS1, BbDXS2*, and *BbDXS3*, annotated as *JF284350, JF284351*and *JF284352*, respectively), and *P. tricornutum DXS* (*Phatr3_Jdraft1689*). Each protein was C-terminally fused with either mVenus or mTurquoise fluorescent proteins. Constructs carrying only promoter, fluorescent proteins and terminator were used as a control.

Diatoms were transformed via bacterial conjugation as described by Diner et al., (2016). To assess the presence and expression of recombinant proteins, transformed algae were cultured in 96-well round-bottom plates (Nunclon-treated, Thermo Fisher Scientific, USA) for 4 days. mVenus fluorescence was measured using a Guava easyCyte flow cytometer (Guava easyCyte HT, Cytek Biosciences, USA) with a 405-nm laser and a 525/40 nm filter. Cultures derived from independent cell lines were analyzed at a flow rate of 0.59 μL/s, collecting 5000 events per sample, in triplicate. Screening for mTurquoise expression was performed using a plate reader set to an excitation wavelength of 434 nm and emission at 474 nm. Fluorescent signals were normalized by calculating the ratio of mTurquoise fluorescence to chlorophyll autofluorescence. From this screening, a minimum of three independent lines exhibiting the highest and the most consistent signal ratios were selected, and the cell line showing the best combination of growth and expression performance was chosen for subsequent experiments

The selected lines were upscaled from 96-well plates, first to 5 mL of ESAW medium and subsequently to 20 mL. This pre-culture was then used to inoculate the experimental culture, which was grown in conditions described in the section “*Algal strains and cultivation,”* in 50 mL of ESAW supplemented with 100 µM Zeocin (Thermo Fisher Scientific, USA).

Consistent phenotypic profiles were observed among independently transformed lines carrying the same construct. This reproducibility supported the use of a single representative line per construct for subsequent analyses and is a key advantage of episomal expression systems. The latter, in fact, enable stable and uniform transgene expression without the variability typically associated with random genomic integration events. Similar consistency in phenotype and expression levels using the same promoter–terminator configuration has been previously demonstrated in *P. tricornutum* episomal systems (Fabris et al., 2020; George et al., 2020).

### Photosynthetic measurements

At specific time points during the experimental period (0, 3, 4, 5, 6 and 7 days after the inoculum), 2 mL of each replicate culture was diluted with 2 mL of ESAW to minimize signal saturation risk for chlorophyll-*a* pulse amplitude modulated (PAM) fluorometry. An AquaPen-C fluorometer (AP 110-C, Photon System Instruments, Brno, Czech Republic) was used for measuring the effective quantum yield (ϕPSII) in light-adapted samples calculated as ΔF/Fm’, where ΔF corresponds to Fm’-F (the maximum minus the minimum fluorescence of light-exposed algae). This measurement was conducted directly on samples obtained from the incubator, without any dark acclimation period to assess the photosynthetic activity under growth light conditions.

Non-photochemical quenching (NPQ) was assessed in transgenic diatom cell lines after 7 days of growth by monitoring changes in the Ft parameter (instantaneous chlorophyll fluorescence) during exposure to high light (HL, 500 μmol photons m⁻² s⁻¹) for 200 seconds, followed by a recovery phase under very low light (LLRec, 10 μmol photons m⁻² s⁻¹) to allow NPQ relaxation (Grouneva et al., 2009). The maximum fluorescence (Fm) was measured by applying a saturating light pulse (SAT) every 20 seconds during both the HL and LLRec phases. NPQ was calculated using the formula (Fm − Fm′) / Fm′, where Fm corresponds to the maximum fluorescence of dark-adapted samples and Fm′ refers to the maximum fluorescence under illuminated conditions, as determined by the SAT pulses.

### Pigment extraction and analysis

Pigment extraction was performed as described by Mendes et al., (2007). Briefly, 5 mL of late-exponential phase diatom culture was centrifuged at 4000 g for 10 min at 21 °C. The pellet was washed with 1.5 mL phosphate-buffered saline (PBS) and centrifuged again under the same conditions. Pellets were flash-frozen in liquid nitrogen and stored at −80 °C until further processing. Samples were freeze-dried overnight, and pigments were extracted using 95% cold methanol buffered with 2% ammonium acetate. Extracts were filtered through 0.2 μm filters, and 10 μL was injected into an Agilent HPLC system equipped with a YMC column (150 × 3.0 mm, S-3 μm). Pigments were eluted with a gradient of solvent A (40% acetone/60% methanol) and solvent B (40% water/60% acetone) as follows: 0–22 min, 0–30% B; 22–41 min, 30–10% B; and 45–55 min, 10–60% B. Column temperature was held at 40 °C with a flow rate of 0.5 mL min⁻¹. Pigments were identified by their absorbance spectra and retention times, compared to pure crystalline standards (DHI, Hørsholm, Denmark).

### Triterpenoid extraction and analysis

Freeze-dried cells from 30 mL of late-exponential phase diatom culture were lysed at 95 °C for 10 min in a 1:1 mixture of 40% KOH and 50% ethanol (250 μL each). Subsequently, 900 μL of hexane was added and the organic phase collected. The extraction was repeated twice, and pooled organic layers were evaporated under nitrogen. For derivatization, 20 μL pyridine and 100 μL N-methyl-N-(trimethylsilyl)trifluoroacetamide were added, followed by incubation at 70 °C for 1 h. Derivatization agents were removed by washing the dried pellets with 100 μL of 50% ethanol, and triterpenoids were extracted via liquid–liquid extraction using 3 volumes of hexane. Extracts were dried and resuspended in 20 μL hexane. GC-FID analysis was performed on an Agilent 7890B using a DB-5 column (30 m × 0.25 mm × 0.25 μm); 1 μL was injected with helium as carrier gas (1 mL/min). Oven program: 80 °C (1 min), ramp to 280 °C at 20 °C/min (hold 45 min), ramp to 320 °C at 20 °C/min (hold 1 min), cool to 80 °C at 50 °C/min. Metabolites were identified by retention times of authentic standards (Merck) and peak areas normalized to the dry weight of the pellets.

### Lipid droplets isolation

Lipid droplets were isolated from algae grown in either nitrate-replete or depleted conditions by adapting the protocol by Yoneda et al. (2016). Briefly, cells were harvested by centrifugation at 3,000 × g for 5 min at room temperature before moving them on ice. The resulting pellet was washed with 10 mL of 10 mM Tris-HCl (pH 7.6) with 2 % (w/v) NaCl and resuspended in 10 of sucrose buffer (0.25 M sucrose, 10 mM Tris-HCl pH 7.6, protease-inhibitor cocktail (Sigma-Aldrich). Cells were disrupted by sonication. Debris and unbroken cells were removed by centrifugation at 50,000 × g for 5 min at 4 °C while the supernatant (containing the crude lipid-droplets fraction) was retained. To generate a discontinuous sucrose gradient, 1 mL of 2.5 M sucrose was added to 4 mL of the supernatant to obtain a final sucrose concentration of 0.7 M. This mixture was transferred to ultracentrifuge tubes and gently overlaid with 5 mL of 10 mM Tris-HCl (pH 7.6). Gradients were then centrifuged at 50,000 g for 20 min at 4 °C, after which the lipid-droplets layer floating at the interface was collected into ten 1.5 mL microcentrifuge tubes. Presence of lipid droplets was confirmed by Nile Red staining. To 200 μL of the lipid droplets isolate, Nile Red was added to a final concentration of 1 μg/mL. Samples were incubated in the dark for 15 minutes at room temperature and subsequently analyzed by confocal microscopy using excitation/emission settings of 488/530 nm. Following staining, the isolated lipids were transferred to a pre-weighed vial, dried, and the vial was reweighed to estimate the total lipid content. The dried lipids were then subjected to squalene extraction using the previously described “triterpenoid extraction” procedure.

### Microscopy

Cells expressing recombinant proteins of interest, at an OD₇₅₀ of 0.2, were used for confocal microscopy analysis. Samples were imaged using a Nikon A1 confocal laser scanning microscope equipped with a Plan Apo λ 100× oil-immersion objective (NA 1.45, Nikon, Japan) at a resolution of 1024 × 512 pixels. mVenus fluorescence was excited with a 488 nm argon laser, and emission was detected between 500–520 nm. mTurquoise was excited with a 405 nm laser, and fluorescence emission was detected between 460–480 nm. Chlorophyll autofluorescence was excited at 561 nm and detected from 625–720 nm. Transmitted light images were acquired using the corresponding DIC channel. All images were processed using ImageJ software.

Coherent Anti-Stokes Raman Scattering (CARS) imaging was performed on a Leica TCS SP8 CARS confocal microscope equipped with a picoEmerald™ picosecond laser system (APE Berlin). The system was tuned to excite the CH₂ symmetric stretching vibration at 2845 cm⁻¹, characteristic of neutral lipids, using a pump beam at 816.4 nm and a Stokes beam at 1064 nm, generating a CARS signal detected at approximately 661 nm. Detection was carried out in the forward or epi-direction using a hybrid detector (HyD) with a 650–700 nm bandpass filter to collect the anti-Stokes signal.

## List of abbreviations

MVA: mevalonate;
MEP: methyl erythritol phosphate;
CARS: Coherent Anti-Stokes Raman Scattering;
PAM: Pulse amplitude modulation;
SQS: squalene synthase;
LD: lipid droplets

## Author contribution (CRediT statement)

LM: Conceptualization, data curation, formal analysis, investigation, methodology, visualization, writing – original draft preparation, review and editing; CJ: investigation, methodology; EAC: methodology, writing – review and editing; MF: Conceptualization, funding acquisition, data curation, methodology, project administration, resources, supervision, writing – original draft preparation (contribution), writing – review and editing.

## Conflict of interests

The authors declare no conflict of interests.

## Acknowledgements

This work was supported by the Novo Nordisk Foundation (grant number NNF22OC0078846 to MF) and by an SDU Climate Cluster (SCC) Research Infrastructure Grant to MF. MF is supported by a Villum Young Investigator Grant (Villum Fonden grant number 37521). The authors would like to thank Lars Duelund for the technical assistance provided with analytical chemistry and the technical team of DaMBIC for the technical assistance provided with analytical chemistry and with microscopy.

## Supporting information

Figures S1–S6 contain additional computational and experimental data including enzyme structure modelling and sequence analyses, engineered strain genotyping, and Table S1-S3 with details on constructs, sequence and primers used in this study.

## Notes

### Competing Interest Statement

The authors have declared no competing interest.

