## Supplementary material for "Iterative and modular expression of *Botryococcus braunii* genes enhances isoprenoid production in the diatom *Phaeodactylum tricornutum*"

### Supporting information file 1

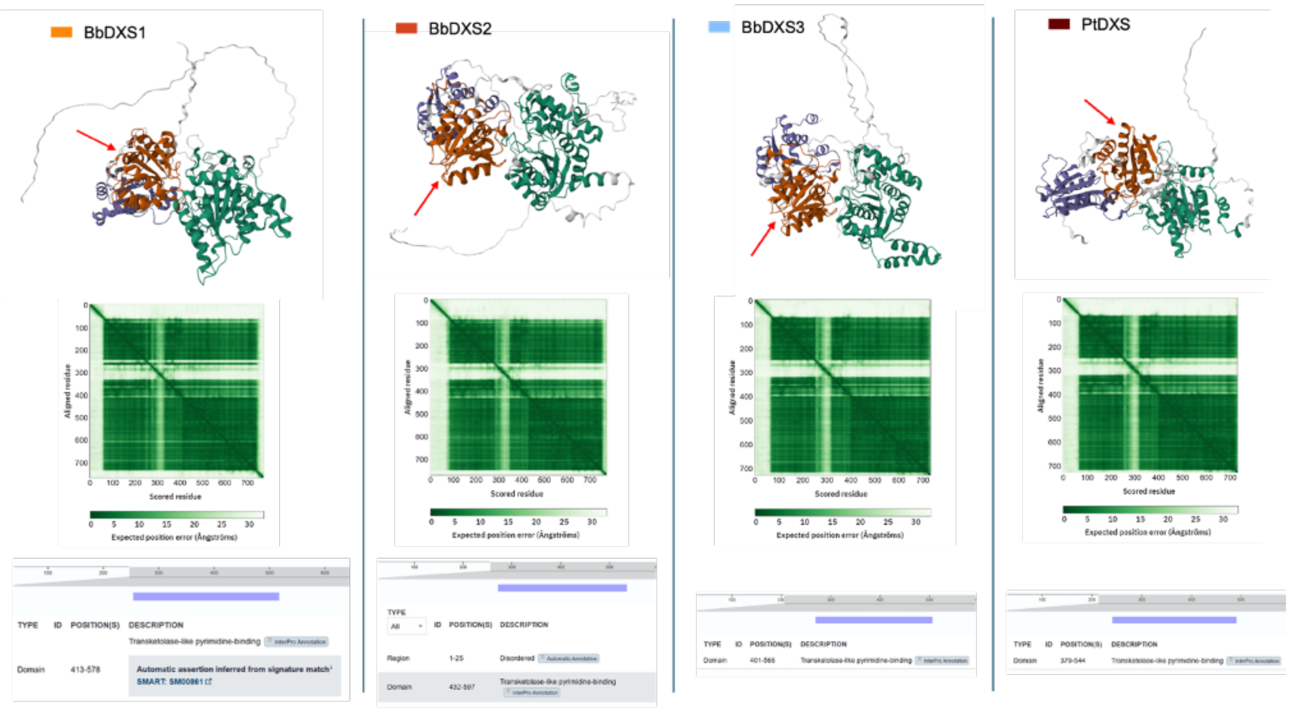


**Figure S1. Structural and domain comparison of *B. braunii* DXS isoforms and *P. tricornutum* DXS.** Predicted tertiary structures of BbDXS1, BbDXS2, BbDXS3, and PtDXS highlight the conserved transketolase-like (pyrimidine-binding) domain (dark orange) and variable N-terminal extensions (gray), with red arrows indicating the putative active site. Contact maps derived from AlphaFold models show conserved folding topology among all isoforms. Bottom panels show domain architecture analyses confirm the presence of the transketolase pyrimidine-binding domain across all sequences.

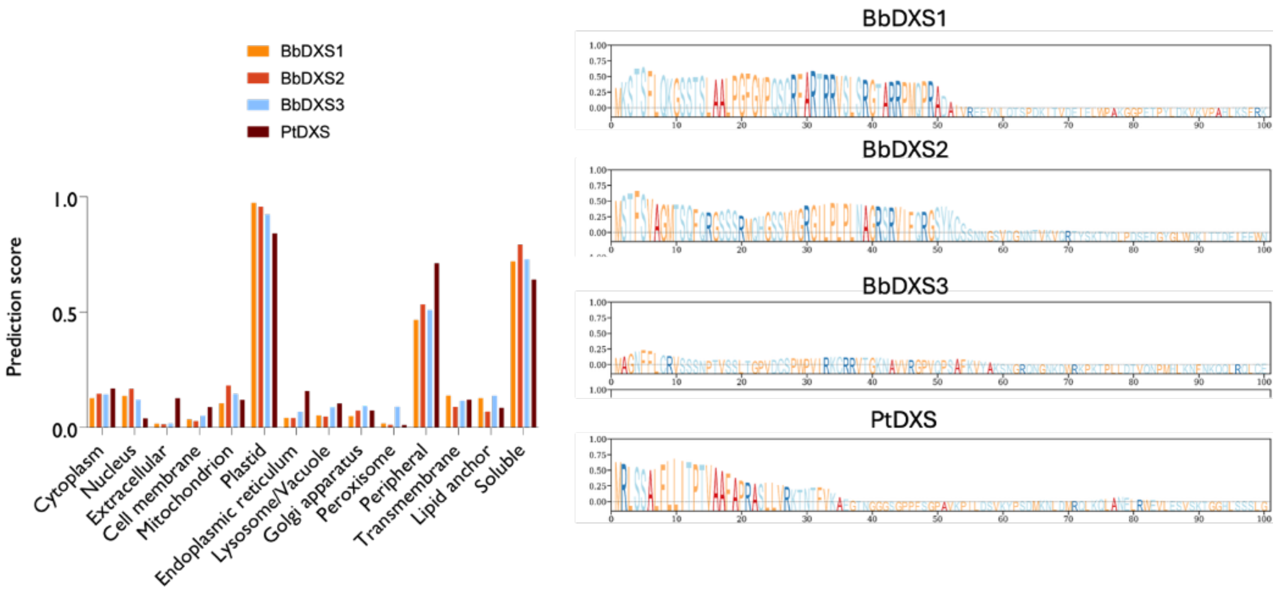


**Figure S2. Amino acid composition and sequence alignment of the N-terminal regions of *B. braunii DXS* isoforms and *P. tricornutum DXS*.** Bar plots show DeepLoc-predicted localization probabilities for BbDXS1, BbDXS2, BbDXS3, and PtDXS across major cellular compartments. Sequence logo plots display the N-terminal regions (first 100 amino acids) of each isoform, illustrating compositional features potentially associated with targeting signals.

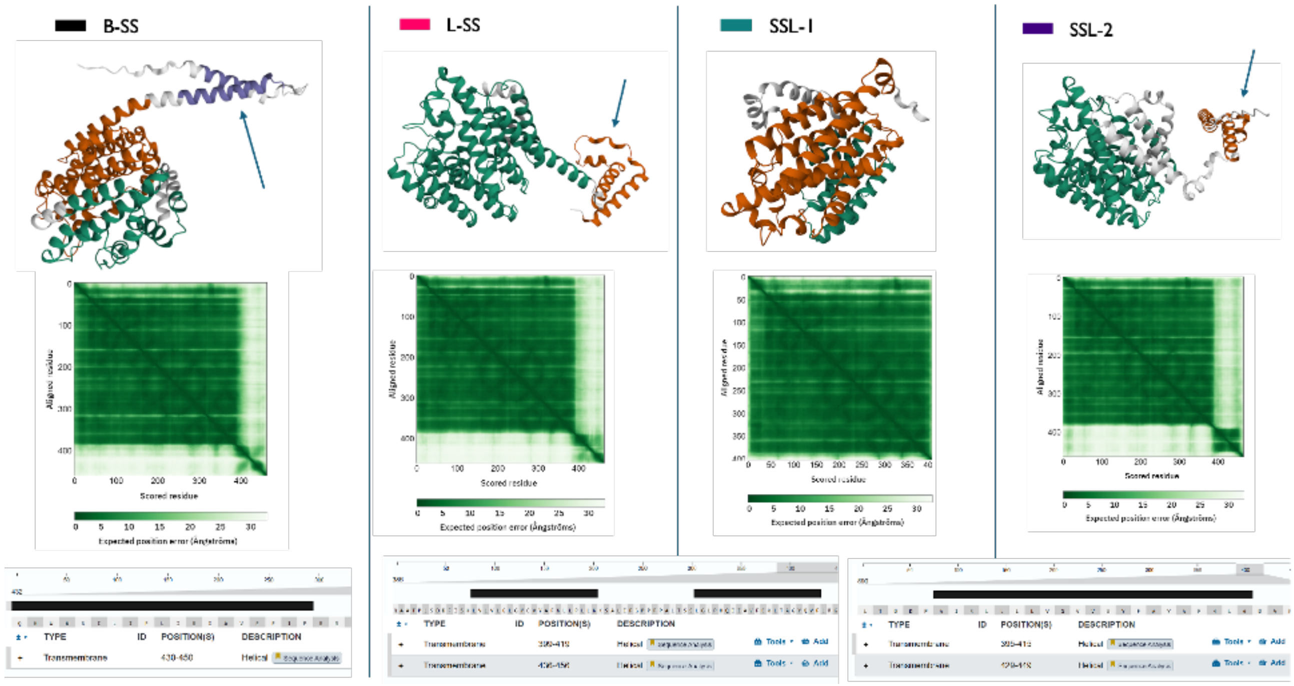


**Figure S3. Structural and domain comparison of *B. braunii*'s SQS isoforms and *P. tricornutum*'s SQS.** Predicted tertiary structures of B-SS, L-SS, SSL-1, and SSL-2 highlight the conserved transmembrane domain (dark orange). Contact maps derived from AlphaFold models show conserved folding topology among all isoforms.

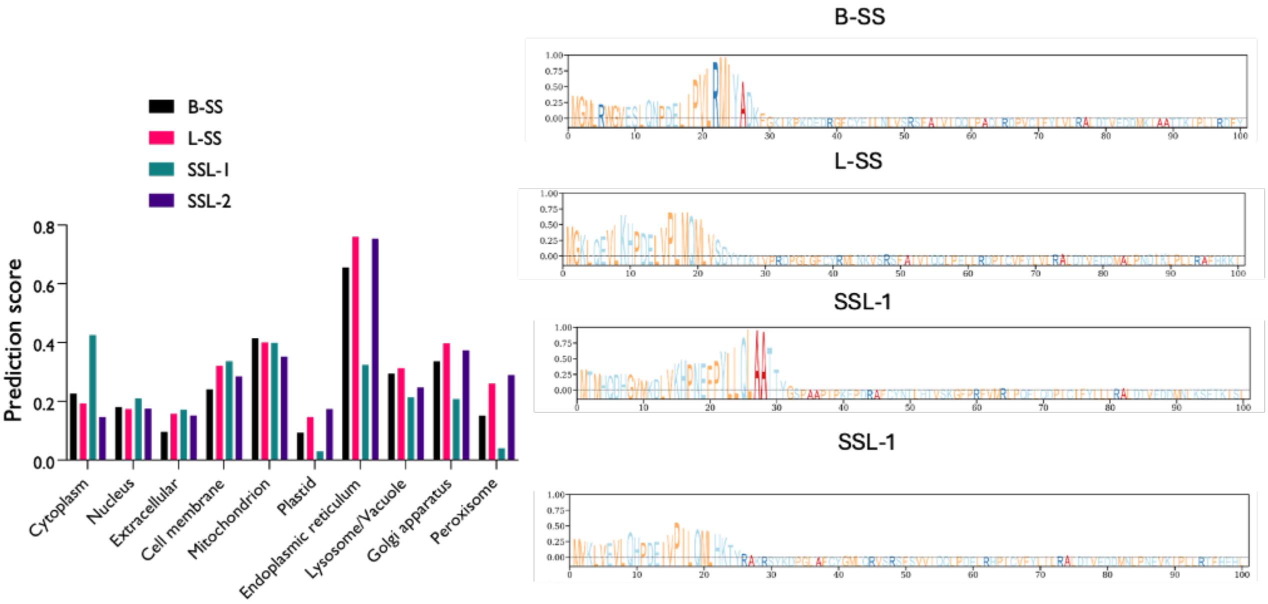


**Figure S4. Amino acid composition and sequence alignment of the N-terminal regions of *B. braunii* *SQS*.** Bar plots show DeepLoc-predicted localization probabilities for B-SS, L-SS, SSL-1, and SSL-2 across major cellular compartments. Sequence logo plots display the N-terminal regions (first 100 amino acids) of each isoform, illustrating compositional features potentially associated with targeting signals.

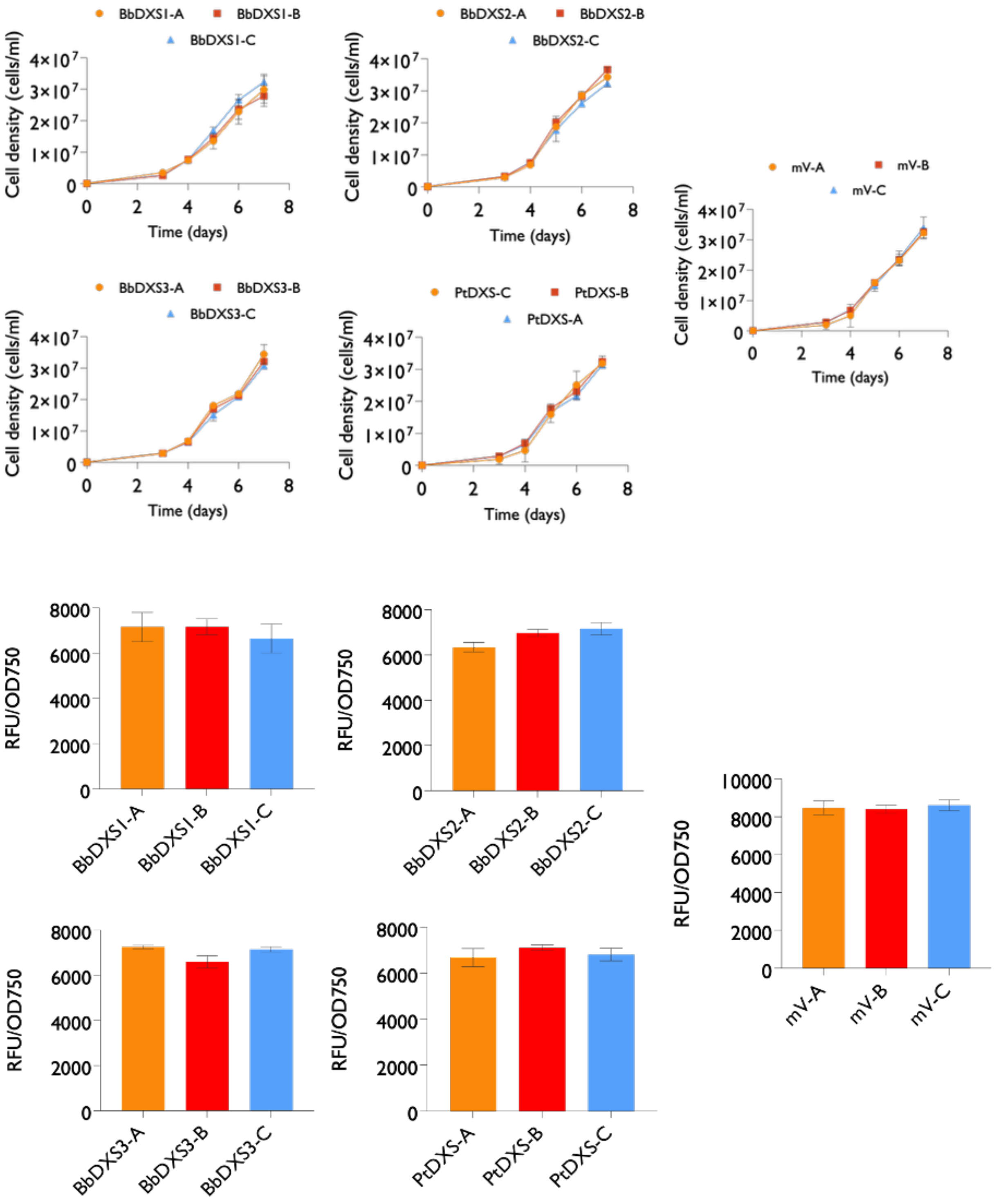


**Figure S5. Growth pattern and recombinant protein fluorescence of three independent cell lines.** Growth curves of transgenic lines overexpressing BbDXS1–3 and PtDXS, expressed as cell density calculated through flow cytometry. Fluorescence of the respective mVenus or mTurquoise fluorescent fusion protein, used as a proxy for recombinant enzyme abundance at harvest and normalized to optical density, measured with a fluorescence plate reader.

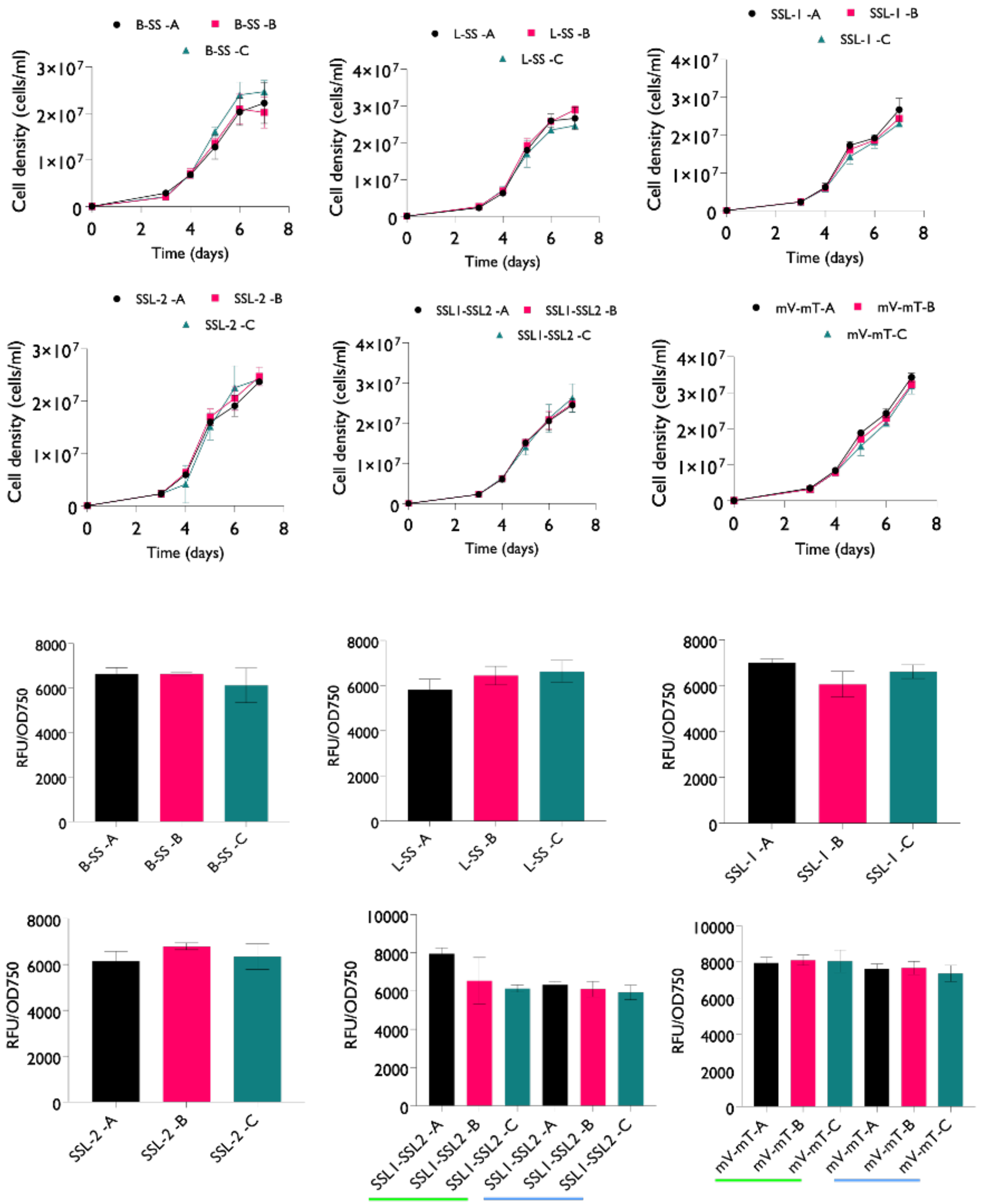


**Figure S5. Growth pattern and fluorescence of three independent cell lines.** Growth curves of transgenic lines overexpressing the different SQS isoforms, expressed as cell density calculated through flow cytometry. Fluorescence of the respective mVenus or mTurquoise fluorescent fusion protein, used as a proxy for recombinant enzyme abundance at harvest and normalized to optical density, measured with a fluorescence plate reader.

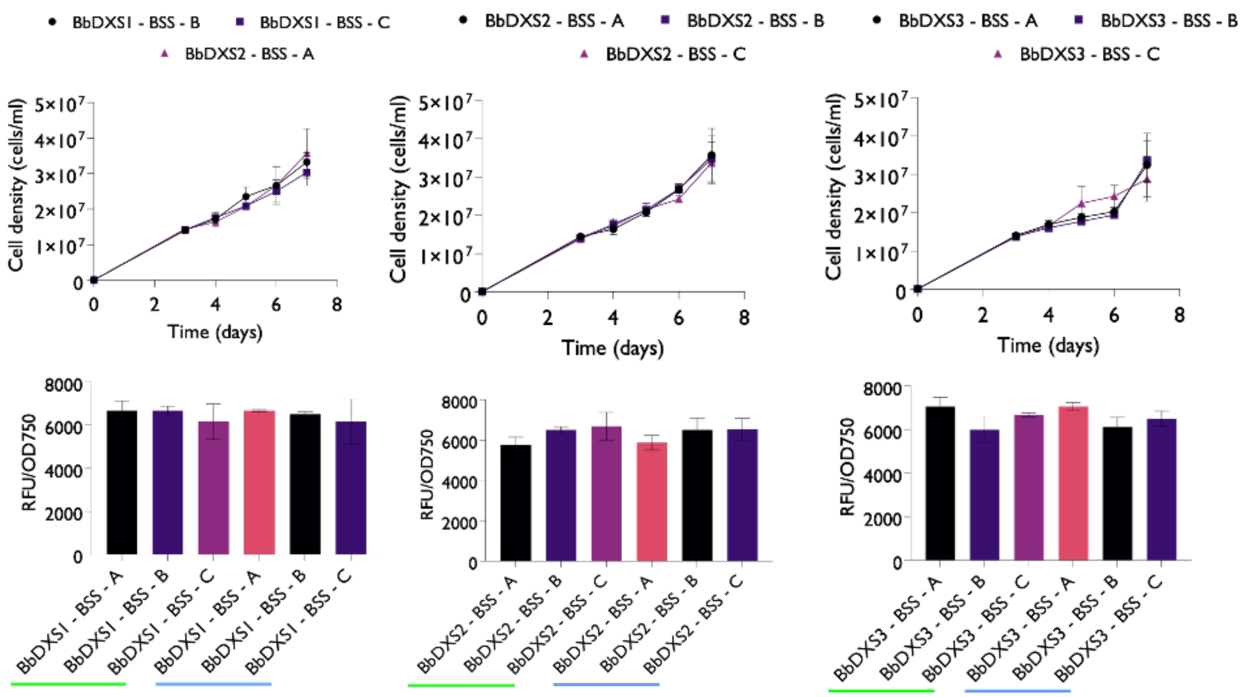


**Figure S6. Growth pattern and fluorescence of three independent cell lines.** Growth curves of transgenic lines overexpressing the combination of the three BbDXS isoforms and B-SS, expressed as cell density calculated through flow cytometry. Fluorescence of the respective mVenus or mTurquoise fluorescent protein, used as a proxy for recombinant enzyme abundance at harvest and normalized to optical density, measured with a fluorescence plate reader.

**Table S1**. L0 parts for construction of episomes using the uLoop assembly method. DNA parts were codon-optimized, synthesized de novo, and obtained from Genewiz (Azenta Life Sciences).

| **L0 part** | **Gene ID / accession** | **Description** | **Reference** |
|---|---|---|---|
| AC_pPt49202 | Phatr3_J49202 | Promoter | (Pollak et al., 2019) |
| L1-1 PTCv2 | N/A | Conjugation and prop. Elements | (Pollak et al., 2019) |
| EF_tPtJ25172 | Phatr3_J25172 | Terminator associated with Phatr3_J25172 promoter (FcbP) | (Pollak et al., 2019) |
| DE_B-SS-mVenus | AF205791 | Squalene biosynthetic pathway gene B-SS fused to fluorescent protein | This study |
| DE_L-SS-mTurquoise | KT388100 | Squalene biosynthetic pathway gene L-SS fused to cyan fluorescent protein | This study |
| CD_SSL1 | HQ585058 | Squalene biosynthetic pathway gene SSL1 | This study |
| CD_SSL2 | G0Y287 | Squalene biosynthetic pathway gene SSL2 | This study |
| CD_BbDXS1 | JF284350 | B. braunii 1-deoxy-D-xylulose-5-phosphate synthase isoform 1 | This study |
| CD_BbDXS2 | JF284351 | B. braunii 1-deoxy-D-xylulose-5-phosphate synthase isoform 2 | This study |
| CD_BbDXS3 | JF284352 | B. braunii 1-deoxy-D-xylulose-5-phosphate synthase isoform 3 | This study |
| CD_PtDXS | Phatr3_Jdraft1689 | P. tricornutum 1-deoxy-D-xylulose-5- | This study |

| | | | |
|---|---|---|---|
| | | phosphate synthase | |
| CD_mVenus | NA | Yellow fluorescent protein | (Pollak et al., 2019) |
| CD_mTurquoise | NA | Turquoise fluorescent protein | (Pollak et al., 2019) |
| DE_VenusThrHISFLAG | NA | Yellow fluorescent protein with a Thormbine, 6xHIS and FLAG (DYKDDDDK) tag JCVI | (Pollak et al., 2019) |
| DE_mTurquoise ThrHISFLAG | NA | Cyan fluorescent protein with a Thormbine, 6xHIS and FLAG (DYKDDDDK) tag JCVI | (Pollak et al., 2019) |
| DE_3xStop | NA | Three stop codon | (Pollak et al., 2019) |

**Table S2. L0 plasmid sequences**

L0_BSS-mVenus_DE 4166 bp

1 cattactcgc atccattctc aggctgtctc gtctcgtctc aggtctccag gtatgggtat
61 gcttcgctgg ggagtcgaat cgttgcagaa tccggatgaa ttaatcccgg tcttgcgtat
121 gatttacgct gataagtttg gaaagatcaa gccaaaggac gaagaccggg gcttctgcta
181 tgaaattctg aaccttgttt caagaagttt tgccatcgtc atccaacaac tccctgcaca
241 gctgagggac ccagtctgca tcttttacct tgtactacgc gccctcgaca ctgtcgaaga
301 tgatatgaaa attgccgcca cgaccaagat tcccttgctg cgtgactttt atgagaaaat
361 ttctgaccgt tccttccgca tgacggccgg agatcaaaag gactacatcc gtctgttgga
421 tcagtacccc aaggtgacaa gcgttttttt gaaattgacc ccccgtgaac aagaaataat
481 tgccgacatt acaaagcgta tgggtaatgg aatggctgac ttcgtgcata agggtgttcc
541 tgacacggtg ggtgactacg acctttactg ccactacgtt gctggggtcg tgggtcttgg
601 gctttcccag ttgttcgttg cgagtggact ccagtcgccc tcgttgaccc gcagtgaaga
661 cctttccaac cacatgggcc tcttccttca gaagaccaac atcatccgcg attactttga
721 ggacattaat gagctccctg ccccccggat gttctggccc cgtgaaatct ggggcaagta
781 tgcgaacaac ctcgctgaat tcaaagaccc ggccaacaag gcggctgcaa tgtgttgcct
841 caacgagatg gtcacagatg cattgcgtca cgcggtgtac tgcctccagt acatgtccat
901 gattgaggat ccgcagatct tcaacttttg tgccatccct cagacgatgg ccttcggcac
961 cctctctttg tgttacaaca actacactat cttcacgggt cccaaggcgg ctgtcaagct
1021 gcgtcgaggt acgactgcca agctgatgta cacctctaac aatatgtttg ccatgtaccg
1081 tcatttcctc aactttgccg agaagctcga agtccgatgt aacaccgaaa cttccgagga
1141 tccgagcgtc accaccactc tggaacacct gcacaagatt aaagctgcct gcaaggctgg
1201 actcgcacgc acaaaagatg acacctttga cgaattgcgc tcccgcttgt tggccctcac
1261 gggaggcagc ttttacctcg cctggaccta caatttccta gatcttcgag gcccgggaga
1321 cttgccaacc ttcctctccg taacccaaca ctggtggtct attctcatct tcctcatttc
1381 gattgccgtc ttctttattc cgtcgcggcc ctcccctcgt cccactctct ccgccatggt
1441 gagcaagggc gaggagctgt tcaccggggt ggtgcccatc ctggtcgagc tggacggcga
1501 cgtaaacggc cacaagttca gcgtgtccgg cgagggcgag ggcgatgcca cctacggcaa
1561 gctgaccctg aagctgatct gcaccaccgg caagctgccc gtgccctggc ccaccctcgt
1621 gaccaccctg ggctacggcc tgcagtgctt cgcccgctac cccgaccaca tgaagcagca
1681 cgacttcttc aagtccgcca tgcccgaagg ctacgtccag gagcgcacca tcttcttcaa
1741 ggacgacggc aactacaaga cccgcgccga ggtgaagttc gagggcgaca ccctggtgaa
1801 ccgcatcgag ctgaagggca tcgacttcaa ggaggacggc aacatcctgg ggcacaagct
1861 ggagtacaac tacaacagcc acaacgtcta tatcaccgcc gacaagcaga agaacggcat
1921 caaggccaac ttcaagatcc gccacaacat cgaggacggc ggcgtgcagc tcgccgacca
1981 ctaccagcag aacaccccca tcggcgacgg ccccgtgctg ctgcccgaca accactacct
2041 gagctaccag tccgccctga gcaaagaccc caacgagaag cgcgatcaca tggtcctgct
2101 ggagttcgtg accgccgccg ggatcactct cggcatggac gagctgtaca agtgataagc
2161 ttcgagacca ccaggataca tagattacca caactccgag cccttccacc cacagaatca
2221 ggggataacg caggaaagaa catgtgagca aaaggccagc aaaaggccag gaaccgtaaa
2281 aaggccgcgt tgctggcgtt tttccatagg ctccgccccc ctgacgagca tcacaaaaat
2341 cgacgctcaa gtcagaggtg gcgaaacccg acaggactat aaagatacca ggcgtttccc
2401 cctggaagct ccctcgtgcg ctctcctgtt ccgaccctgc cgcttaccgg atacctgtcc
2461 gcctttctcc cttcgggaag cgtggcgctt tctcatagct cacgctgtag gtatctcagt
2521 tcggtgtagg tcgttcgctc caagctgggc tgtgtgcacg aaccccccgt tcagcccgac
2581 cgctgcgcct tatccggtaa ctatcgtctt gagtccaacc cggtaagaca cgacttatcg
2641 ccactggcag cagccactgg taacaggatt agcagagcga ggtatgtagg cggtgctaca
2701 gagttcttga agtggtggcc taactacggc tacactagaa gaacagtatt tggtatctgc
2761 gctctgctga agccagttac cttcggaaaa agagttggta gctcttgatc cggcaaacaa
2821 accaccgctg gtagcggtgg tttttttgtt tgcaagcagc agattacgcg cagaaaaaaa

2881 ggatctcaag aagatccttt gatcttttct acggggtctg acgctcagtg gaacgaaaac
2941 tcacgttaag ggattttggt catgcattct aggtgattat tatttgccga ctaccttggt
3001 gatctcgcct ttcacgtagt ggacaaattc ttccaactga tctgcgcgcg aggccaagcg
3061 atcttcttct tgtccaagat aagcctgtct agcttcaagt atgacgggct gatactgggc
3121 cggcaggcgc tccattgccc agtcggcagc gacatccttc ggcgcgattt tgccggttac
3181 tgcgctgtac caaatgcggg acaacgtaag cactacattt cgctcatcgc cagcccagtc
3241 gggcggcgag ttccatagcg ttaaggtttc atttagcgcc tcaaatagat cctgttcagg
3301 aaccggatca aagagttcct ccgccgctgg acctaccaag gcaacgctat gttctcttgc
3361 ttttgtcagc aagatagcca gatcaatgtc gatcgtggct ggctcgaaga taccagcaag
3421 aatgtcattg cgctgccatt ctccaaattg cagttcgcgc ttagctggat aacgccacgg
3481 aatgatgtcg tcgtgcacaa caatggtgac ttctacagcg cggagaatct cactctctcc
3541 aggggaagcc gaagtttcca aaaggtcgtt gatcaaagct cgccgcgttg tttcatcaag
3601 ccttacggtc accgtaacca gcaaatcaat atcactgtgt ggcttcaggc cgccatccac
3661 tgcggagccg tacaaatgta cggccagcaa cgtcggttcg agatggcgct cgatgacgcc
3721 aactacctct gatagttgag tcgatacttc ggcgatcacc gcttccctca tgcgaaacga
3781 tcctcatcct gtctcttgat cagatattga tcccctgcgc catcagatcc ttggcggcaa
3841 gaaagccatc cagtttactt tgcagggctt cccaacctta ccagagggcg ccccagctgg
3901 caattccggt tcgcttgcta agcttgcatg cctgcaagtc gactctagag gagcggtgat
3961 cacaggcagc aacgctctgt catcgttaca atcaacatgc taccctccgc gagatcatcc
4021 gtgtttcaaa cccggcagct tagttgccgt tcttccgaat agcatcggta acatgagcaa
4081 agtctgccgc cttacaacgg ctctcccgct gacgccgtcc cggactgatg ggctgcctgt
4141 atcgagtggt gattttgtgc cgagct

L0_LSS-Turquoise_DE        4169 bp

1 cattactcgc atccattctc aggctgtctc gtctcgtctc aggtctccag gtatggggaa
61 gctacaagaa gttttgaagc atccagacga attggtccct cttatgcaga tgctcgtctc
121 cgactactac acaaagatcg tcccgcgtga ccccggtcta ggcttttgct atcgtatgtt
181 gaataaagtt agtcgcagtt ttgcaattgt catccaacag cttcctgaac ttctccggga
241 tccaatttgc gtgttctacc ttgtacttcg cgcccttgac actgttgaag atgatatggc
301 tctcccaaat gatatcaagc ttccgctcct ccgtgcattc cataagaaga tctatgaccg
361 aaaatggtct atgaagtgtg gttacggacc ctatgtgcaa ctgatggagg aatatccgat
421 ggtgacaggt gtctttctca agttggaccc cggcccgcgt gaggtgatta cggaaatttg
481 tcgaaagatg ggagcgggaa tggccgaatt cattcccaaa gaagtactga ctgtgaagga
541 ctacgaccag tactgccact acgctgccgg actcgtggga gagggtttgt ccaagcttgc
601 cgtcggttcg ggattggaaa accccgtcct cttgcaaaag gaggacctct caaatcacat
661 gggcctgttt ctgcagaaga ccaacatcgt ccgtgattac ttggaagata tcaacgaaga
721 acccgcgccc cggatgttct ggcccaagga aatttgggga aaatacacca aggacttggc
781 tgattttaag gatccggcca atgagaaggg tgcggtccag tgcttgaacc acatggtcac
841 ggacgctctg cgacacgggg agcacgcgct gaagtacatg gccctgctgc gcgacccgca
901 atactttaac ttctgcgcca tccctcaagt catggccttc ggtaccttat cgctctgcta
961 caacaacccc caggtcttta aaggtgttgt gaaattgcgt aagggcgaga gcgccaagct
1021 gatgactacg gtcaaatcga tgccggctct ataccgcacc ttcttgcgta tggccgatga
1081 catggtggcc cgttgtaagg gagaagctcg tcaggatcca aacgtcgcca ccaccctcaa
1141 gcgcctccaa gccattcagg cggtttgtaa gaccggcctc agatcttcca tcaaatctcg
1201 caagaagcag gccgccactc cgctcagcga cgacttcatt tccaagttgg tcttggtact
1261 tggtctggga tactgcgttt acgcatttaa cctcctcccg ctcttgtgga aatcggccct
1321 cattcccggc cctccccctc ccgccctcac ctcctccttg ggcctccctc accagattat
1381 cgctgttttc tgtgttctca cggctggcta ccaggtcttc cttcgcggag gccttgccat
1441 ggtgagcaag ggcgaggagc tgttcaccgg ggtggtgccc atcctggtcg agctggacgg
1501 cgacgtaaac ggccacaagt tcagcgtgtc cggcgagggc gagggcgatg ccacctacgg

1561 caagctgacc ctgaagttca tctgcaccac cggcaagctg cccgtgccct ggcccaccct
1621 cgtgaccacc ctgtcctggg gcgtgcagtg cttcgcccgc taccccgacc acatgaagca
1681 gcacgacttc ttcaagtccg ccatgcccga aggctacgtc caggagcgca ccatcttctt
1741 caaggacgac ggcaactaca agacccgcgc cgaggtgaag ttcgagggcg acaccctggt
1801 gaaccgcatc gagctgaagg gcatcgactt caaggaggac ggcaacatcc tggggcacaa
1861 gctggagtac aactacttta gcgacaacgt ctatatcacc gccgacaagc agaagaacgg
1921 catcaaggcc aacttcaaga tccgccacaa catcgaggac ggcggcgtgc agctcgccga
1981 ccactaccag cagaacaccc ccatcggcga cggccccgtg ctgctgcccg acaaccacta
2041 cctgagcacc cagtccaagc tgagcaaaga ccccaacgag aagcgcgatc acatggtcct
2101 gctggagttc gtgaccgccg ccgggatcac tctcggcatg gacgagctgt acaagtgata
2161 agcttcgaga ccaccaggat acatagatta ccacaactcc gagcccttcc acccacagaa
2221 tcaggggata acgcaggaaa gaacatgtga gcaaaaggcc agcaaaaggc caggaaccgt
2281 aaaaaggccg cgttgctggc gtttttccat aggctccgcc cccctgacga gcatcacaaa
2341 aatcgacgct caagtcagag gtggcgaaac ccgacaggac tataaagata ccaggcgttt
2401 ccccctggaa gctccctcgt gcgctctcct gttccgaccc tgccgcttac cggatacctg
2461 tccgcctttc tcccttcggg aagcgtggcg ctttctcata gctcacgctg taggtatctc
2521 agttcggtgt aggtcgttcg ctccaagctg ggctgtgtgc acgaaccccc cgttcagccc
2581 gaccgctgcg ccttatccgg taactatcgt cttgagtcca acccggtaag acacgactta
2641 tcgccactgg cagcagccac tggtaacagg attagcagag cgaggtatgt aggcggtgct
2701 acagagttct tgaagtggtg gcctaactac ggctacacta gaagaacagt atttggtatc
2761 tgcgctctgc tgaagccagt taccttcgga aaaagagttg gtagctcttg atccggcaaa
2821 caaaccaccg ctggtagcgg tggttttttt gtttgcaagc agcagattac gcgcagaaaa
2881 aaaggatctc aagaagatcc tttgatcttt tctacggggt ctgacgctca gtggaacgaa
2941 aactcacgtt aagggatttt ggtcatgcat tctaggtgat tattatttgc cgactacctt
3001 ggtgatctcg cctttcacgt agtggacaaa ttcttccaac tgatctgcgc gcgaggccaa
3061 gcgatcttct tcttgtccaa gataagcctg tctagcttca agtatgacgg gctgatactg
3121 ggccggcagg cgctccattg cccagtcggc agcgacatcc ttcggcgcga ttttgccggt
3181 tactgcgctg taccaaatgc gggacaacgt aagcactaca tttcgctcat cgccagccca
3241 gtcgggcggc gagttccata gcgttaaggt ttcatttagc gcctcaaata gatcctgttc
3301 aggaaccgga tcaaagagtt cctccgccgc tggacctacc aaggcaacgc tatgttctct
3361 tgcttttgtc agcaagatag ccagatcaat gtcgatcgtg gctggctcga agataccagc
3421 aagaatgtca ttgcgctgcc attctccaaa ttgcagttcg cgcttagctg gataacgcca
3481 cggaatgatg tcgtcgtgca caacaatggt gacttctaca gcgcggagaa tctcactctc
3541 tccaggggaa gccgaagttt ccaaaaggtc gttgatcaaa gctcgccgcg ttgtttcatc
3601 aagccttacg gtcaccgtaa ccagcaaatc aatatcactg tgtggcttca ggccgccatc
3661 cactgcggag ccgtacaaat gtacggccag caacgtcggt tcgagatggc gctcgatgac
3721 gccaactacc tctgatagtt gagtcgatac ttcggcgatc accgcttccc tcatgcgaaa
3781 cgatcctcat cctgtctctt gatcagatat tgatcccctg cgccatcaga tccttggcgg
3841 caagaaagcc atccagttta ctttgcaggg cttcccaacc ttaccagagg gcgccccagc
3901 tggcaattcc ggttcgcttg ctaagcttgc atgcctgcaa gtcgactcta gaggagcggt
3961 gatcacaggc agcaacgctc tgtcatcgtt acaatcaaca tgctaccctc cgcgagatca
4021 tccgtgtttc aaacccggca gcttagttgc cgttcttccg aatagcatcg gtaacatgag
4081 caaagtctgc cgccttacaa cggctctccc gctgacgccg tcccggactg atgggctgcc
4141 tgtatcgagt ggtgattttg tgccgagct

CD_L0_B-BbSSL1 3268 bp DNA

1 cattactcgc atccattctc aggctgtctc gtctcgtctc aggtctccaa tgacgatgca
61 ccaggaccac ggagtcatga aagaccttgt caagcatcca aatgaatttc catacttgct
121 ccaactcgcg gccacaacgt acggttcgcc ggctgccccg atcccgaagg aaccggaccg
181 agcgttctgc tacaacactc ttcacaccgt ttcgaagggt ttccccagat ttgttatgcg

241 ccttccgcag gaactccaag atccgatatg catcttctac ctcttgttgc gagctctaga
301 cacggtggaa gacgacatga acctcaagag tgaaacgaag atttcactcc tacgcgtttt
361 ccatgaacac tgttcggacc gtaactggag catgaaatcg gattacggta tctatgccga
421 tcttatggaa cgtttcccct tggtcgtatc cgtcttagag aagctccctc ccgccaccca
481 gcagactttc cgtgaaaatg tcaaatacat gggcaatgga atggcagatt ttattgataa
541 gcagatcctg acagtggatg agtacgacct ctactgtcac tatgtggccg gcagttgcgg
601 cattgctgtc accaaggtca ttgtgcagtt caaccttgcc acgcctgaag ctgactccta
661 cgacttttcc aacagcctgg gactcttgct tcagaaggcc aacatcatca ccgactacaa
721 cgaagacatc aacgaagaac ctcgtccccg aatgttctgg ccccaggaga tttgggggaa
781 gtacgcggag aagttggctg actttaatga acccgaaaat attgataccg ccgtgaagtg
841 cttgaaccac atggtcaccg atgcaatgcg tcacattgag ccttccctca agggcatggt
901 ttatttcacc gacaagacag tctttcgggc gctcgctctt ttgctggtca cagcctttgg
961 tcatttgtcc accttgtaca acaacccgaa tgtctttaaa gagaaagttc gtcagcggaa
1021 gggacgcatt gcccgcctgg tcatgagttc ccgtaacgta cctggactct tccgtacctg
1081 tctcaaactc gccaacaact ttgaatcccg ctgcaagcaa gagacggcaa acgatcccac
1141 tgtggccatg actatcaagc gcttgcaatc tattcaagct acctgtcgtg atggactggc
1201 caagtacgac accccctctg gtctgaaatc tttctgcgcc gccccaactc ccaccaagtc
1261 aggtcgagac caccaggata catagattac cacaactccg agcccttcca cccacagaat
1321 caggggataa cgcaggaaag aacatgtgag caaaaggcca gcaaaaggcc aggaaccgta
1381 aaaaggccgc gttgctggcg tttttccata ggctccgccc ccctgacgag catcacaaaa
1441 atcgacgctc aagtcagagg tggcgaaacc cgacaggact ataaagatac caggcgtttc
1501 cccctggaag ctccctcgtg cgctctcctg ttccgaccct gccgcttacc ggatacctgt
1561 ccgcctttct cccttcggga agcgtggcgc tttctcatag ctcacgctgt aggtatctca
1621 gttcggtgta ggtcgttcgc tccaagctgg gctgtgtgca cgaacccccc gttcagcccg
1681 accgctgcgc cttatccggt aactatcgtc ttgagtccaa cccggtaaga cacgacttat
1741 cgccactggc agcagccact ggtaacagga ttagcagagc gaggtatgta ggcggtgcta
1801 cagagttctt gaagtggtgg cctaactacg gctacactag aagaacagta tttggtatct
1861 gcgctctgct gaagccagtt accttcggaa aaagagttgg tagctcttga tccggcaaac
1921 aaaccaccgc tggtagcggt ggtttttttg tttgcaagca gcagattacg cgcagaaaaa
1981 aaggatctca agaagatcct ttgatctttt ctacggggtc tgacgctcag tggaacgaaa
2041 actcacgtta agggattttg gtcatgcatt ctaggtgatt attatttgcc gactaccttg
2101 gtgatctcgc ctttcacgta gtggacaaat tcttccaact gatctgcgcg cgaggccaag
2161 cgatcttctt cttgtccaag ataagcctgt ctagcttcaa gtatgacggg ctgatactgg
2221 gccggcaggc gctccattgc ccagtcggca gcgacatcct tcggcgcgat tttgccggtt
2281 actgcgctgt accaaatgcg ggacaacgta agcactacat ttcgctcatc gccagcccag
2341 tcgggcggcg agttccatag cgttaaggtt tcatttagcg cctcaaatag atcctgttca
2401 ggaaccggat caaagagttc ctccgccgct ggacctacca aggcaacgct atgttctctt
2461 gcttttgtca gcaagatagc cagatcaatg tcgatcgtgg ctggctcgaa gataccagca
2521 agaatgtcat tgcgctgcca ttctccaaat tgcagttcgc gcttagctgg ataacgccac
2581 ggaatgatgt cgtcgtgcac aacaatggtg acttctacag cgcggagaat ctcactctct
2641 ccaggggaag ccgaagtttc caaaaggtcg ttgatcaaag ctcgccgcgt tgtttcatca
2701 agccttacgg tcaccgtaac cagcaaatca atatcactgt gtggcttcag gccgccatcc
2761 actgcggagc cgtacaaatg tacggccagc aacgtcggtt cgagatggcg ctcgatgacg
2821 ccaactacct ctgatagttg agtcgatact tcggcgatca ccgcttccct catgcgaaac
2881 gatcctcatc ctgtctcttg atcagatatt gatcccctgc gccatcagat ccttggcggc
2941 aagaaagcca tccagtttac tttgcagggc ttcccaacct taccagaggg cgccccagct
3001 ggcaattccg gttcgcttgc taagcttgca tgcctgcaag tcgactctag aggagcggtg
3061 atcacaggca gcaacgctct gtcatcgtta caatcaacat gctaccctcc gcgagatcat
3121 ccgtgtttca aacccggcag cttagttgcc gttcttccga atagcatcgg taacatgagc
3181 aaagtctgcc gccttacaac ggctctcccg ctgacgccgt cccggactga tgggctgcct
3241 gtatcgagtg gtgattttgt gccgagct

CD_L0_B-BbSSL2_ 3454 bp DNA circular UNA 09-NOV-2023

1 cattactcgc atccattctc aggctgtctc gtctcgtctc aggtctccaa tggtcaaact
61 cgtcgaagtt ttgcagcacc cggacgaaat cgtccccatc ctgcagatgt tgcacaagac
121 ctaccgcgca aagcgcagct acaaagaccc tggtctagcc ttttgctacg gaatgttgca
181 acgtgtttcg agaagctttt cagttgttat ccagcagctc cctgacgaat tgcgccaccc
241 gatttgcgtg ttttacctta ttcttcgtgc cctcgatact gtcgaagatg acatgaacct
301 cccaaacgaa gttaagattc ctcttcttcg taccttccat gaacatctct ttgaccggtc
361 gtggaagctc aaatgtggat acggaccgta tgtagatttg atggaaaact atcccctagt
421 cacggatgtc ttccttacac tctcgcccgg cgcacaagag gtaatccggg actcgacgcg
481 ccgcatgggt aatggtatgg ccgacttcat tggcaaggat gaggtccact ctgtcgcgga
541 gtacgatctg tactgtcact atgtggctgg tttggtcggt tccgctgtgg ccaagatttt
601 tgttgacagc gggctggaaa aggaaaatct cgtcgcggag gtggatctgg ccaacaacat
661 gggccagttc ctccaaaaga ccaacgttat tcgagactac ttggaggata ttaatgaaga
721 accggcccct cgtatgttct ggccgcgtga aatttggggc aaatacgccc aggaacttgc
781 ggactttaag gacccagcca acgaaaaagc ggccgtacag tgcctgaatc acatggtcac
841 tgatgcactc cgacactgcg aaatcggcct taacgtgatt ccgctgttgc agaacattgg
901 catcctccgc tcatgcctca tccccgaagt catgggattg cgtaccctca ccttgtgtta
961 caacaatccc caagtcttcc gtggcgtggt caagatgcgt cgtggggaaa ctgccaagtt
1021 gttcatgagt atctacgaca agcgctcctt ctaccaaacc tacctccgac tcgccaacga
1081 gttggaagca aagtgtaaag gagaggcgag tggagatccc atggttgcca cgacgctcaa
1141 gcatgtgcac ggaatccaga agtcctgcaa agccgctctc tcctccaagg agctgcttgc
1201 caagtctggt tcggccctca cagacgatcc cgctatccgt ttgctgcttc tcgtcggagt
1261 cgttgcctac tttgcctacg ccttcaactt gggagacgtc cggggagagc acggtgtccg
1321 tgctctcggt tccattttgg acctatccca aaagggcttg gctgtggcga gtgtcgctct
1381 gcttctctta gtgcttctgg ccaggtcccg ccttcccttg ctcacctctg cttcttccaa
1441 gcagtcaggt cgagaccacc aggatacata gattaccaca actccgagcc cttccaccca
1501 cagaatcagg ggataacgca ggaaagaaca tgtgagcaaa aggccagcaa aaggccagga
1561 accgtaaaaa ggccgcgttg ctggcgtttt tccataggct ccgcccccct gacgagcatc
1621 acaaaaatcg acgctcaagt cagaggtggc gaaacccgac aggactataa agataccagg
1681 cgtttccccc tggaagctcc ctcgtgcgct ctcctgttcc gaccctgccg cttaccggat
1741 acctgtccgc ctttctccct tcgggaagcg tggcgctttc tcatagctca cgctgtaggt
1801 atctcagttc ggtgtaggtc gttcgctcca agctgggctg tgtgcacgaa ccccccgttc
1861 agcccgaccg ctgcgcctta tccggtaact atcgtcttga gtccaacccg gtaagacacg
1921 acttatcgcc actggcagca gccactggta acaggattag cagagcgagg tatgtaggcg
1981 gtgctacaga gttcttgaag tggtggccta actacggcta cactagaaga acagtatttg
2041 gtatctgcgc tctgctgaag ccagttacct tcggaaaaag agttggtagc tcttgatccg
2101 gcaaacaaac caccgctggt agcggtggtt tttttgtttg caagcagcag attacgcgca
2161 gaaaaaaagg atctcaagaa gatcctttga tcttttctac ggggtctgac gctcagtgga
2221 acgaaaactc acgttaaggg attttggtca tgcattctag gtgattatta tttgccgact
2281 accttggtga tctcgccttt cacgtagtgg acaaattctt ccaactgatc tgcgcgcgag
2341 gccaagcgat cttcttcttg tccaagataa gcctgtctag cttcaagtat gacgggctga
2401 tactgggccg gcaggcgctc cattgcccag tcggcagcga catccttcgg cgcgattttg
2461 ccggttactg cgctgtacca aatgcgggac aacgtaagca ctacatttcg ctcatcgcca
2521 gcccagtcgg gcggcgagtt ccatagcgtt aaggtttcat ttagcgcctc aaatagatcc
2581 tgttcaggaa ccggatcaaa gagttcctcc gccgctggac ctaccaaggc aacgctatgt
2641 tctcttgctt ttgtcagcaa gatagccaga tcaatgtcga tcgtggctgg ctcgaagata
2701 ccagcaagaa tgtcattgcg ctgccattct ccaaattgca gttcgcgctt agctggataa
2761 cgccacggaa tgatgtcgtc gtgcacaaca atggtgactt ctacagcgcg gagaatctca
2821 ctctctccag gggaagccga agtttccaaa aggtcgttga tcaaagctcg ccgcgttgtt

2881 tcatcaagcc ttacggtcac cgtaaccagc aaatcaatat cactgtgtgg cttcaggccg
2941 ccatccactg cggagccgta caaatgtacg gccagcaacg tcggttcgag atggcgctcg
3001 atgacgccaa ctacctctga tagttgagtc gatacttcgg cgatcaccgc ttccctcatg
3061 cgaaacgatc ctcatcctgt ctcttgatca gatattgatc ccctgcgcca tcagatcctt
3121 ggcggcaaga aagccatcca gtttactttg cagggcttcc caaccttacc agagggcgcc
3181 ccagctggca attccggttc gcttgctaag cttgcatgcc tgcaagtcga ctctagagga
3241 gcggtgatca caggcagcaa cgctctgtca tcgttacaat caacatgcta ccctccgcga
3301 gatcatccgt gtttcaaacc cggcagctta gttgccgttc ttccgaatag catcggtaac
3361 atgagcaaag tctgccgcct tacaacggct ctcccgctga cgccgtcccg gactgatggg
3421 ctgcctgtat cgagtggtga ttttgtgccg agct

L0_-_BbDXS1_CD 4369 bp

1 cattactcgc atccattctc aggctgtctc gtctcgtctc aggtctccaa tgaagtccac
61 gtcttttctt caaaagggct caagcacaag cctcgctgca cttccaggtt tcggagtccc
121 gcaatcctgt cgttttgccc gtacgcggcg agtttccctt tcgcgcggaa ccgcccgtcg
181 ccctatgcag ccacgcgcgg acgcaatcgt cagagaagaa gttaacttgc agaccagccc
241 ggataagata accgtggatg agattgaact ctggcccgcc aagggtggac cagaaacgcc
301 ataccttgac aaagtcaagg tccctgcgca cttgaagtcc ttccgtaagg atgagttgaa
361 gacagtctgc cgtgagctcc gcgctgaaat catcaacgcc gtgtctgcca cgggaggaca
421 cttgggttcc tccctgggag tagtggagct tactgttgct atccactacg tcttcgactg
481 cccggaggac aagcttgttt gggacgtggg ccaccaggca tacggtcaca aaattttgac
541 tgggcgtagg gacaaaatgc acaccattcg acaacgagac ggcttgtctg gattcaccaa
601 ccgctcggaa agtgaatacg acgcctttgg tgccggtcac agttcgacgt ccatctcggc
661 tgccctcgga atggcggtcg gacgtgatct tctgggaaag gataaccact gcattgccgt
721 cattggtgat ggagccatta ctggtggtat ggcgtacgag gccctcaacc atgctggctt
781 cctggggtcc gaaaaaatcc aaggaaaggg gaacaacatc ggccgcacga ttgtcatcct
841 caatgacaac cagcaggtca gtctcccaac ccaattcaat ggagagaaac aaaagcctgt
901 tggggctttg gctgacgccc tctctcaagt cgcatctaaa caactttcca actccgccaa
961 agaaattgtg aagcagctgc cggagccctt gcaggctgtt ggaggccagt tggaaaaggt
1021 tgtctctaca atcggtggaa ataacacctt ctttgacgag ttgggtgtgg cccatgtagg
1081 ccctattgat ggccacaacg tggaagacct cgtgaacgtc ttggaatgga ttaagtcaca
1141 gaaggacgga ggtcccgtca ttgtccatat cttgacagaa aaaggatatg gatatgagtt
1201 tgctgaaaag gcctcggacc gtatgcatgg agtcgcgaag tatgacgtca cctctggcaa
1261 gcaagtcaag agttccagca aggtggcctc ctacaccact tactttgcag actcattaat
1321 cgcggaagcc gaacgtgacg gccgggtcat tggtatccat gccgccatgg gtggtggtac
1381 cggcatgaac cgctttgcca agcgattccc caagcgcacc tttgatgttg gcattgctga
1441 acagcacgcg gtcaccttcg ccgccggcct ggcctgtgaa ggacttatcc cgatgtgcgc
1501 aatttactcc tccttcctac agcgtgccta cgaccaggtc atccatgacg ttgccctcca
1561 aaacctcccg gtgcgctttg ccatggatcg cgcgggcctc gttggggcgg atggggctac
1621 gcacagtgga tttgcagacg tcacctacat ggcatgtgtc cccaacatga tcgtaatggc
1681 cccctcgaac gaagccgagc tgtgtaatgc cgtggccact tccattgcca tcgactttgc
1741 tccgtcgtgc ttccggttcc cgcgtggtaa tggaattggc gtcgacctgg ctgaatacgg
1801 tgtgcaaccc aacttcaagg gcaccccttg ggagattggt aaaggcaaaa tccgccggaa
1861 cggatcgaac aataagccgg aaggagatgt agctcttttg ggttacggca cggtggtcaa
1921 tgattgcctc gccgcggccg agatgctaga agcccaagga attaagacca ccgtagctga
1981 tatgcgtttc tgcaagcctt tggatgaaga actcattgtg cagctggcca agaaccaccc
2041 ggtcgtcatt actgttgaag aaaacactgt tggtggcttt gcctcccacg tcctgcactt
2101 tatgacagag cagggcttcc tcgatggaaa gatgaagctc cgtacccttt tgctccccga
2161 tcgtttcatt gaacacggca cccagcagca gcagctagaa gaagccctcc tcaccgctaa
2221 tgacattgta gagacagcgg tcaatgcttt gggtattgct cccgttcctc aaatcatcat

2281 cccccctcag caggttgtta cgcccccccca ggccgtcagc ccgccccagg tcgccactcc
2341 cacgcccgcc cccctcaact caggtcgaga ccaccaggat acatagatta ccacaactcc
2401 gagcccttcc acccacagaa tcaggggata acgcaggaaa gaacatgtga gcaaaaggcc
2461 agcaaaaggc caggaaccgt aaaaaggccg cgttgctggc gtttttccat aggctccgcc
2521 cccctgacga gcatcacaaa aatcgacgct caagtcagag gtggcgaaac ccgacaggac
2581 tataaagata ccaggcgttt ccccctggaa gctccctcgt gcgctctcct gttccgaccc
2641 tgccgcttac cggatacctg tccgcctttc tcccttcggg aagcgtggcg ctttctcata
2701 gctcacgctg taggtatctc agttcggtgt aggtcgttcg ctccaagctg ggctgtgtgc
2761 acgaaccccc cgttcagccc gaccgctgcg ccttatccgg taactatcgt cttgagtcca
2821 acccggtaag acacgactta tcgccactgg cagcagccac tggtaacagg attagcagag
2881 cgaggtatgt aggcggtgct acagagttct tgaagtggtg gcctaactac ggctacacta
2941 gaagaacagt atttggtatc tgcgctctgc tgaagccagt taccttcgga aaaagagttg
3001 gtagctcttg atccggcaaa caaaccaccg ctggtagcgg tggttttttt gtttgcaagc
3061 agcagattac gcgcagaaaa aaaggatctc aagaagatcc tttgatcttt tctacggggt
3121 ctgacgctca gtggaacgaa aactcacgtt aagggatttt ggtcatgcat tctaggtgat
3181 tattatttgc cgactacctt ggtgatctcg cctttcacgt agtggacaaa ttcttccaac
3241 tgatctgcgc gcgaggccaa gcgatcttct tcttgtccaa gataagcctg tctagcttca
3301 agtatgacgg gctgatactg ggccggcagg cgctccattg cccagtcggc agcgacatcc
3361 ttcggcgcga ttttgccggt tactgcgctg taccaaatgc gggacaacgt aagcactaca
3421 tttcgctcat cgccagccca gtcgggcggc gagttccata gcgttaaggt ttcatttagc
3481 gcctcaaata gatcctgttc aggaaccgga tcaaagagtt cctccgccgc tggacctacc
3541 aaggcaacgc tatgttctct tgcttttgtc agcaagatag ccagatcaat gtcgatcgtg
3601 gctggctcga agataccagc aagaatgtca ttgcgctgcc attctccaaa ttgcagttcg
3661 cgcttagctg gataacgcca cggaatgatg tcgtcgtgca caacaatggt gacttctaca
3721 gcgcggagaa tctcactctc tccaggggaa gccgaagttt ccaaaaggtc gttgatcaaa
3781 gctcgccgcg ttgtttcatc aagccttacg gtcaccgtaa ccagcaaatc aatatcactg
3841 tgtggcttca ggccgccatc cactgcggag ccgtacaaat gtacggccag caacgtcggt
3901 tcgagatggc gctcgatgac gccaactacc tctgatagtt gagtcgatac ttcggcgatc
3961 accgcttccc tcatgcgaaa cgatcctcat cctgtctctt gatcagatat tgatcccctg
4021 cgccatcaga tccttggcgg caagaaagcc atccagttta ctttgcaggg cttcccaacc
4081 ttaccagagg gcgccccagc tggcaattcc ggttcgcttg ctaagcttgc atgcctgcaa
4141 gtcgactcta gaggagcggt gatcacaggc agcaacgctc tgtcatcgtt acaatcaaca
4201 tgctaccctc cgcgagatca tccgtgtttc aaacccggca gcttagttgc cgttcttccg
4261 aatagcatcg gtaacatgag caaagtctgc cgccttacaa cggctctccc gctgacgccg
4321 tcccggactg atgggctgcc tgtatcgagt ggtgattttg tgccgagct

L0_-_BbDXS2_CD 4372 bp DNA

1 cattactcgc atccattctc aggctgtctc gtctcgtctc aggtctccaa tgtcgacgtt
61 ttcggtcgct ggcatgacgt cccagttcca acgtggatcc agctcccgta tgcagcatgg
121 ctcgtctgtc gtcggacgcg gtattcttcc gctgccccta aatgctggac gctctcgtgt
181 aatctttcag agaggatctt acaagtgctc atcaaacaat ggaagtgtag acggtaacaa
241 caccgttaag gtacagcgaa cctacagtaa gacatacgac ctccccgact ccgaagatgg
301 gtacggacta tgggacaaga tcaccaccga cgagattgaa gaatggaaca atggtccaga
361 gactcccctc ctcgacacca ttaaatcccc gctgggcttg aaaaactaca gcatgccgga
421 acttcgtgcc ctctgcaaag aattacgcgc tgacatcatc cacaccgtct cgcaaaccgg
481 agggcacctg ggatcatccc tcggtgtggc tgaattgacc gtggcccttc attatgtctt
541 caactgtcct tacgacaaaa ttatttggga cgtcggccat caggcctacg gccacaagat
601 cttgaccggc cggagggccc agatgcccac cattcgccag tacaagggct tgtctggttt
661 cacaaagcgt tccgagagcg agtacgacgc ctttggtgcc ggccacagct ccacttccat
721 atcggccgct ctgggaatgg ccgttggtcg tgatttcaag aaccaaaaca accacgccat

781 tgcggtcatt ggagatggag ctattaccgg tggcatggcc tacgaggccc tcaaccatgc
841 cggtttcatg aaaacggaca aaatgattgt cattctcaac gacaataagc aggtttctct
901 cccgacgcag tacaatgccg gagatcaaaa gccggtggga gccttggccg ctaccttggc
961 cgaaatccaa gctgacccga acctccggga aatccgtgaa acgattaaga actttgctga
1021 acagatgcct gcccccatcc gcgattttgc caaggccgtg gatgatgtcg ggcacgacat
1081 tttgtctcaa aatagtggag ccaaattgtt caaggatttg aagctctggt acatcggacc
1141 cgtggacggt cacgatgtag gccttcttgt caaggtcctg caggatgtga agaaacgtcc
1201 ggacatgggc cctgtcctca ttcacatcct cacggagaag ggccgtggat atgagcccgc
1261 cgaaacttct atggatcgta tgcacggcgt tggaaagtac gacacgaaga ccggtaaggc
1321 gctccccagt acctccaaga gtaagagcta cacaaacgtc tacgcggact cgctcgcagc
1381 agaagctgcg cttgacccca aaatcttggc cgtccatgcc gccatgggcg gaggaactgg
1441 tatgaaccga ttcgaaaagc gcttcgcttc gcgcactttt gatgttggta tcgccgaaca
1501 gcacgcggtc acttttgctg ctggactcgc atcggagggt ttgaaaccct tctgtgccat
1561 ttactccacg ttcctccagc gcgccttcga ccagatcatc cacgacgttg cactccagaa
1621 tctgccagtc cgttttgcca ttgaccgtgg aggccttgtc ggagcggacg gcgccacgca
1681 cggtggattt gccgatgtga cctacatggc atgcgtgcca aacatggttg tcatggcgcc
1741 atcgaacgaa gccgagttgt gccacgccgt cgccaccgcc gccgcgtata acgatggacc
1801 ctcctgtttc cgcttccccc gcggtgacgg tatcggtgtc gacatggaag ctgaaggtgt
1861 cgaccctaag tccttcaagg gtcaaccttg ggagatcggt aagggagtta ttcgacggct
1921 tggcaagcac gcggccctct tgggctacgg cacaatggtg aacgattgtt tggccgctgc
1981 agaagaacta gctaagaaag gaattgatgt tgccgtaatg gatatgcgat tttgcaagcc
2041 tctggacaca caactcgttg agcaagccgc gaagaactat ccgatcgtta ttactgtgga
2101 ggaaaatgct atcggcggtt ttggctccca cgttgccaac cacctggtga agaccggtct
2161 tctggatacg gcggtgaagt tccgttccat gatcctcccg gaccgctaca ttgatcacgg
2221 aacacaggcg caacagcgtg aagaagccgg cttgaatacc actaccattg tggctaaggt
2281 cgaaagtctt atgccctaca actccaacaa cggggtccat gcaaatggtg tccacgcaaa
2341 cggggtccat gccaatgggg tgtcaggtcg agaccaccag gatacataga ttaccacaac
2401 tccgagccct tccacccaca gaatcagggg ataacgcagg aaagaacatg tgagcaaaag
2461 gccagcaaaa ggccaggaac cgtaaaaagg ccgcgttgct ggcgtttttc cataggctcc
2521 gcccccctga cgagcatcac aaaaatcgac gctcaagtca gaggtggcga aacccgacag
2581 gactataaag ataccaggcg tttccccctg gaagctccct cgtgcgctct cctgttccga
2641 ccctgccgct taccggatac ctgtccgcct ttctcccttc gggaagcgtg gcgctttctc
2701 atagctcacg ctgtaggtat ctcagttcgg tgtaggtcgt tcgctccaag ctgggctgtg
2761 tgcacgaacc ccccgttcag cccgaccgct gcgccttatc cggtaactat cgtcttgagt
2821 ccaacccggt aagacacgac ttatcgccac tggcagcagc cactggtaac aggattagca
2881 gagcgaggta tgtaggcggt gctacagagt tcttgaagtg gtggcctaac tacggctaca
2941 ctagaagaac agtatttggt atctgcgctc tgctgaagcc agttaccttc ggaaaaagag
3001 ttggtagctc ttgatccggc aaacaaacca ccgctggtag cggtggtttt tttgtttgca
3061 agcagcagat tacgcgcaga aaaaaaggat ctcaagaaga tcctttgatc ttttctacgg
3121 ggtctgacgc tcagtggaac gaaaactcac gttaagggat tttggtcatg cattctaggt
3181 gattattatt tgccgactac cttggtgatc tcgcctttca cgtagtggac aaattcttcc
3241 aactgatctg cgcgcgaggc caagcgatct tcttcttgtc caagataagc ctgtctagct
3301 tcaagtatga cgggctgata ctgggccggc aggcgctcca ttgcccagtc ggcagcgaca
3361 tccttcggcg cgattttgcc ggttactgcg ctgtaccaaa tgcgggacaa cgtaagcact
3421 acatttcgct catcgccagc ccagtcgggc ggcgagttcc atagcgttaa ggtttcattt
3481 agcgcctcaa atagatcctg ttcaggaacc ggatcaaaga gttcctccgc cgctggacct
3541 accaaggcaa cgctatgttc tcttgctttt gtcagcaaga tagccagatc aatgtcgatc
3601 gtggctggct cgaagatacc agcaagaatg tcattgcgct gccattctcc aaattgcagt
3661 tcgcgcttag ctggataacg ccacggaatg atgtcgtcgt gcacaacaat ggtgacttct
3721 acagcgcgga gaatctcact ctctccaggg gaagccgaag tttccaaaag gtcgttgatc
3781 aaagctcgcc gcgttgtttc atcaagcctt acggtcaccg taaccagcaa atcaatatca

3841 ctgtgtggct tcaggccgcc atccactgcg gagccgtaca aatgtacggc cagcaacgtc
3901 ggttcgagat ggcgctcgat gacgccaact acctctgata gttgagtcga tacttcggcg
3961 atcaccgctt ccctcatgcg aaacgatcct catcctgtct cttgatcaga tattgatccc
4021 ctgcgccatc agatccttgg cggcaagaaa gccatccagt ttactttgca gggcttccca
4081 accttaccag agggcgcccc agctggcaat tccggttcgc ttgctaagct tgcatgcctg
4141 caagtcgact ctagaggagc ggtgatcaca ggcagcaacg ctctgtcatc gttacaatca
4201 acatgctacc ctccgcgaga tcatccgtgt ttcaaacccg gcagcttagt tgccgttctt
4261 ccgaatagca tcggtaacat gagcaaagtc tgccgcctta caacggctct cccgctgacg
4321 ccgtcccgga ctgatgggct gcctgtatcg agtggtgatt ttgtgccgag ct

LOCUS L0_-_BbDXS3_CD 4249 bp

1 cattactcgc atccattctc aggctgtctc gtctcgtctc aggtctccaa tggctggcaa
61 cttcttcctt tgtcgtgtgt cttcgtcaaa ccccaccgtt agttctctga caggccctgt
121 ggactgctcc ccctggcccg tcattcgcaa gtgccgtcgc gttactggaa agaacgccgt
181 agtacgcgga cccgtccaac ctagtgcctt caaggtttac gccaagtcca atggtcgtca
241 aaacggtaac aaagactggc gtaaacccaa aaccccccctc ctggatacgg tccagaatcc
301 catgcacctc aagaatttca acaagcaaca gctccgtcag ctgtgtgagg agcttcgtca
361 agacatcatt agcagtgttt ccaccaccgg aggtcacttg ggatcgtctc tgggtgttgt
421 cgaattgacg gtggcccttc actacgtctt caacgctcca tacgatcgta ttatttggga
481 cgtcggccac caggcttacg gacacaagat cctcaccggg cgtagggacc gcatgaatac
541 cattcgacag tctggaggcc tttcggggtt caccaaccga agcgaaagcg actacgatgc
601 ctttggtgca ggccatagct cgacttccat ctcagcggct gtaggaatgg ctattggatt
661 ggatatcaag ggcatgagaa accactccat tgctgtcatt ggggatggag ctattacagg
721 aggcatggcc tacgaggcta tgaaccacgc cggattcgag aacgtcaaga acctcatcat
781 catcctcaat gataatcagc aggtatctct ccccacgcac tacaatggac agcaacagtc
841 gcctgtcggc gccctgtctg ccaccttggc taagctccag accaacaagg gtctgctaga
901 actccgggag cgtgtcaaag ccttcactaa gactctgccg gcaccgctcc aagacgccac
961 cgccaagatg gacgagtacg cccgtggcat gatttccggt gtggcatcca ccttgttcga
1021 ggaactcggc tgcttctatg tcggtccgat tgacggtcac gacgtgagtc agctcgtcga
1081 cattctgcga gacttaaagg aaaccccagt ccagggaccg gtcttgcttc atgtgcggac
1141 gcagaagggc cagggctaca agtttgctga agaagcctcg gaccgcatgc atggcgtggg
1201 caagtacgac cccagaaactg gtgaacaatt caaaggtgct gagaagacgc cgagttacac
1261 acatttcttt gcggatgcgt tgattgccga agccgaactt gactcccgtg ttcaaggaat
1321 ccacgccgct atgggtggtg gtaccggcat gaaccgcttt gcgaagcgtt tccaatcccg
1381 gacatatgac gttggcatcg ctgaacagca cgcggtgacc tttgcagccg gactcgcctg
1441 cgaaggattg gttccggtcg ttgccatcta ctcctccttt ctccaacgtg gctatgacca
1501 agtcatccat gacgtagcgc tccagtccct tcccgtccgc tttgcaatgg accgggccgg
1561 acttgtgggg gccgatggag ccactcactc cggttttgcg gacgtcacct tcatgggctg
1621 cctccccaac atggtggtca tggcgccatc gaacgaagcc gaactgtgca acgccctcgc
1681 cacggcgatc gcatacgatg aaggtccctc gtgtttccgt tttccccgcg gtaacggcat
1741 tggcgttgat ttggcggaat ttggagtcgc ccctaactac aagggtcgcc cttgggtcct
1801 tgggaaggcc tcaattcgcc gaagcgggaa ggatgttgcc tttttggctt acggaaccgt
1861 tgtcaatgac tgccttgccg ccgccgagat gctcgcggcc aacggagtct ccgccaccgt
1921 ggtagatatg cgcttctgta agcctctcga taaagaaatc gtgaaacgct gtgccaaaga
1981 acacccggtc gtcattacag tcgaagaaaa tgccatcggt ggttttggat cccacgttct
2041 tcactgcttg gctttggagg gattgctgga caccaacgtc aaggtccgtc ccatggtgct
2101 cccggatcgt ttcattgagc atgccacgca gcaggaacag ctcgctacgg ctggtctaga
2161 tgcggctcac atatttgcca ctgccatcaa gacgttggga ttgccttaca acgccccggc
2221 agtcctagcc ggtgcattgt caggtcgaga ccaccaggat acatagatta ccacaactcc
2281 gagcccttcc acccacagaa tcaggggata acgcaggaaa gaacatgtga gcaaaaggcc

2341 agcaaaaggc caggaaccgt aaaaaggccg cgttgctggc gtttttccat aggctccgcc
2401 cccctgacga gcatcacaaa aatcgacgct caagtcagag gtggcgaaac ccgacaggac
2461 tataaagata ccaggcgttt ccccctggaa gctccctcgt gcgctctcct gttccgaccc
2521 tgccgcttac cggatacctg tccgcctttc tcccttcggg aagcgtggcg ctttctcata
2581 gctcacgctg taggtatctc agttcggtgt aggtcgttcg ctccaagctg ggctgtgtgc
2641 acgaaccccc cgttcagccc gaccgctgcg ccttatccgg taactatcgt cttgagtcca
2701 acccggtaag acacgactta tcgccactgg cagcagccac tggtaacagg attagcagag
2761 cgaggtatgt aggcggtgct acagagttct tgaagtggtg gcctaactac ggctacacta
2821 gaagaacagt atttggtatc tgcgctctgc tgaagccagt taccttcgga aaaagagttg
2881 gtagctcttg atccggcaaa caaaccaccg ctggtagcgg tggttttttt gtttgcaagc
2941 agcagattac gcgcagaaaa aaaggatctc aagaagatcc tttgatcttt tctacggggt
3001 ctgacgctca gtggaacgaa aactcacgtt aagggatttt ggtcatgcat tctaggtgat
3061 tattatttgc cgactacctt ggtgatctcg cctttcacgt agtggacaaa ttcttccaac
3121 tgatctgcgc gcgaggccaa gcgatcttct tcttgtccaa gataagcctg tctagcttca
3181 agtatgacgg gctgatactg ggccggcagg cgctccattg cccagtcggc agcgacatcc
3241 ttcggcgcga ttttgccggt tactgcgctg taccaaatgc gggacaacgt aagcactaca
3301 tttcgctcat cgccagccca gtcgggcggc gagttccata gcgttaaggt ttcatttagc
3361 gcctcaaata gatcctgttc aggaaccgga tcaaagagtt cctccgccgc tggacctacc
3421 aaggcaacgc tatgttctct tgcttttgtc agcaagatag ccagatcaat gtcgatcgtg
3481 gctggctcga agataccagc aagaatgtca ttgcgctgcc attctccaaa ttgcagttcg
3541 cgcttagctg gataacgcca cggaatgatg tcgtcgtgca caacaatggt gacttctaca
3601 gcgcggagaa tctcactctc tccaggggaa gccgaagttt ccaaaaggtc gttgatcaaa
3661 gctcgccgcg ttgtttcatc aagccttacg gtcaccgtaa ccagcaaatc aatatcactg
3721 tgtggcttca ggccgccatc cactgcggag ccgtacaaat gtacggccag caacgtcggt
3781 tcgagatggc gctcgatgac gccaactacc tctgatagtt gagtcgatac ttcggcgatc
3841 accgcttccc tcatgcgaaa cgatcctcat cctgtctctt gatcagatat tgatcccctg
3901 cgccatcaga tccttggcgg caagaaagcc atccagttta ctttgcaggg cttcccaacc
3961 ttaccagagg gcgccccagc tggcaattcc ggttcgcttg ctaagcttgc atgcctgcaa
4021 gtcgactcta gaggagcggt gatcacaggc agcaacgctc tgtcatcgtt acaatcaaca
4081 tgctaccctc cgcgagatca tccgtgtttc aaacccggca gcttagttgc cgttcttccg
4141 aatagcatcg gtaacatgag caaagtctgc cgccttacaa cggctctccc gctgacgccg
4201 tcccggactg atgggctgcc tgtatcgagt ggtgattttg tgccgagct
